# Movement and performance predict widespread cortical activity in a visual detection task

**DOI:** 10.1101/709642

**Authors:** David B. Salkoff, Edward Zagha, Erin McCarthy, David A. McCormick

## Abstract

Recent studies in mice reveal widespread cortical signals during task performance, however the various task-related and task-independent processes underlying this activity are incompletely understood. Here we recorded wide-field neural activity, as revealed by GCaMP6s, from dorsal cortex while simultaneously monitoring orofacial movements, walking, and arousal (pupil diameter) of head-fixed mice performing a Go/NoGo visual detection task and examined the ability of task performance and spontaneous or task-related movements to predict cortical activity. A linear model was able to explain a significant fraction (33-55% of variance) of widefield dorsal cortical activity, with the largest factors being movements (facial, walk, eye), response choice (hit, miss, false alarm), and arousal, and indicate that a significant fraction of trial-to-trial variability arises from both spontaneous and task-related changes in state (e.g. movements, arousal). Importantly, secondary motor cortex was highly correlated with lick rate, critical for optimal task performance (high d’), and was the first region to significantly predict the lick response on target trials. These findings suggest that secondary motor cortex is critically involved in the decision and performance of learned movements and indicate that a significant fraction of trial-to-trial variation in cortical activity results from spontaneous and task-related movements and variations in behavioral/arousal state.

## Introduction

Complex learned behavior requires transformations of sensory representations into motor outputs via interactions with contextual information. In cerebral cortex, these transformations evolve over time and space as information is passed from sensory to motor regions (de Lafuente and Romo 2006; Siegel, Buschman, and Miller 2015; Guo et al. 2014). How sensory and motor signals are represented within cortex, how these signals propagate between cortical regions, and how activity in different cortical regions relate to specific features of task performance are active areas of research.

Foundational studies of decision-making in the cerebral cortex have identified specialized cortical regions with sensory, motor, or decision-related processing (Georgopoulos, Kettner, and Schwartz 1988; Robinson, Bowman, and Kertzman 1995; Britten et al. 1996; Bisley and Goldberg 2003; Erlich, Bialek, and Brody 2011; Komiyama et al. 2010; Hanks and Summerfield 2017). These studies used focal electrophysiological or imaging recordings that necessarily restricted investigation to one or a few cortical areas. Recent advancements in the development of fluorescence-based neural activity sensors and widefield or two-photon imaging have now made it possible to measure population activity from large regions of cortex simultaneously in awake animals at moderate-to-high temporal resolution (up to 30 Hz with calcium sensors or 500 Hz with voltage sensors). These tools enable the monitoring of decision-related processing across cortex.

Large-scale imaging of dorsal cortex has recently been applied to mice performing a diversity of sensory-motor tasks (Goard et al. 2016; Wekselblatt et al. 2016; Allen et al. 2017; Kyriakatos et al. 2016; Makino et al. 2017; Stringer et al. 2019; Musall et al. 2019). An unexpected finding throughout these studies is the widespread activation of cortex during task performance. One possible explanation is that task-specific sensory, decision and motor processing is broadly distributed across multiple cortical regions (Hernández et al. 2010). A second possibility is that these widespread signals are not directly related to decision-making per se, but can be accounted for by behavioral measures such as movements of the face and body (e.g. whisking, eye movements, locomotion) and arousal (Musall et al. 2019; Stringer et al. 2019; McGinley et al. 2015).

Mice frequently engage in spontaneous behaviors (e.g. movements, changes in arousal) that alter brain dynamics and sensory evoked activity (McGinley et al. 2015; Stringer et al. 2019; Musall et al. 2019; Drew, Winder, and Zhang 2018). Furthermore, the post-stimulus behavioral response of rodents is difficult to control, and responses are sometimes much more complicated than necessary for reward (Kawai et al. 2015). Thus, not monitoring behavioral and arousal state may severely impair one’s ability to account for changes in neural activity during task performance and potentially confound the interpretation of neural signals related to decision-making with changes in state.

Here we recorded wide-field calcium activity from the dorsal cerebral cortex of mice performing a visual detection task. We observed widespread activity shortly after target presentation, with broad regions of parietal and frontal cortex predicting the decision of the mouse to respond (lick). A region of the secondary motor cortex appears to be critically involved in generating appropriate responses, since activity in this region is highly correlated with response initiation, it displays early-onset of choice-related activity, and suppression of this region impairs optimal task performance (see also (Guo et al. 2014; Li et al. 2016; Allen et al. 2017)). Additionally, we monitored facial movements, locomotion, and pupil diameter as measures of behavioral and arousal state. We used a linear model with these measures and task features as regressors to deconstruct the widespread cortical activity into its component processes. We found that a large fraction of cortical activity was related to spontaneous and task-aligned movements of the face, eyes, and body, and that these measures accounted for a significant fraction of trial-to-trial variability of cortical activity (see also (Musall et al. 2019; Stringer et al. 2019)). Our results demonstrate a mixture of task-specific and task-unrelated processes that contribute to neural activity during task performance. These findings highlight the importance of careful behavioral monitoring when relating neural activity to decision-making.

## Materials and Methods

### Animal Subjects

Adult male and female Snap25-2A-GCaMP6s-D mice (2-3 months age; Jackson Labs) were housed with *ad libitum* access to food but were water restricted once behavioral training began. Mice received a minimum of 1 ml per day but could receive more during the task. Body weight was measured multiple times weekly to ensure mice stayed above 85% baseline. Mice were singly housed on a standard light cycle. Experiments were run during the day. All experimental procedures were approved by the Yale Institutional Animal Care & Use Committee.

### Stimuli

The visual target was either a higher-intensity or lower-intensity LED flash presented pseudo-randomly (three trials of each within each block were permuted). In the head-fixed setup, the flash (50 ms) was a dim point of blue light 9 cm from the eye, 80 degrees from directly ahead of the mouse. Intensity was adjusted per day and mouse so that the lower-intensity stimulus elicited 50-60% hit rates (Fig. 1C). The lowest intensity presented was 0.04 µW (for the higher intensity stimulus) and 0.01 µW (for the lower intensity stimulus). The auditory distractor stimulus was a 50 ms sweep from 4-16 kHz, with a sound pressure level of 65 dB at the location of the mouse.

**Figure 1.**
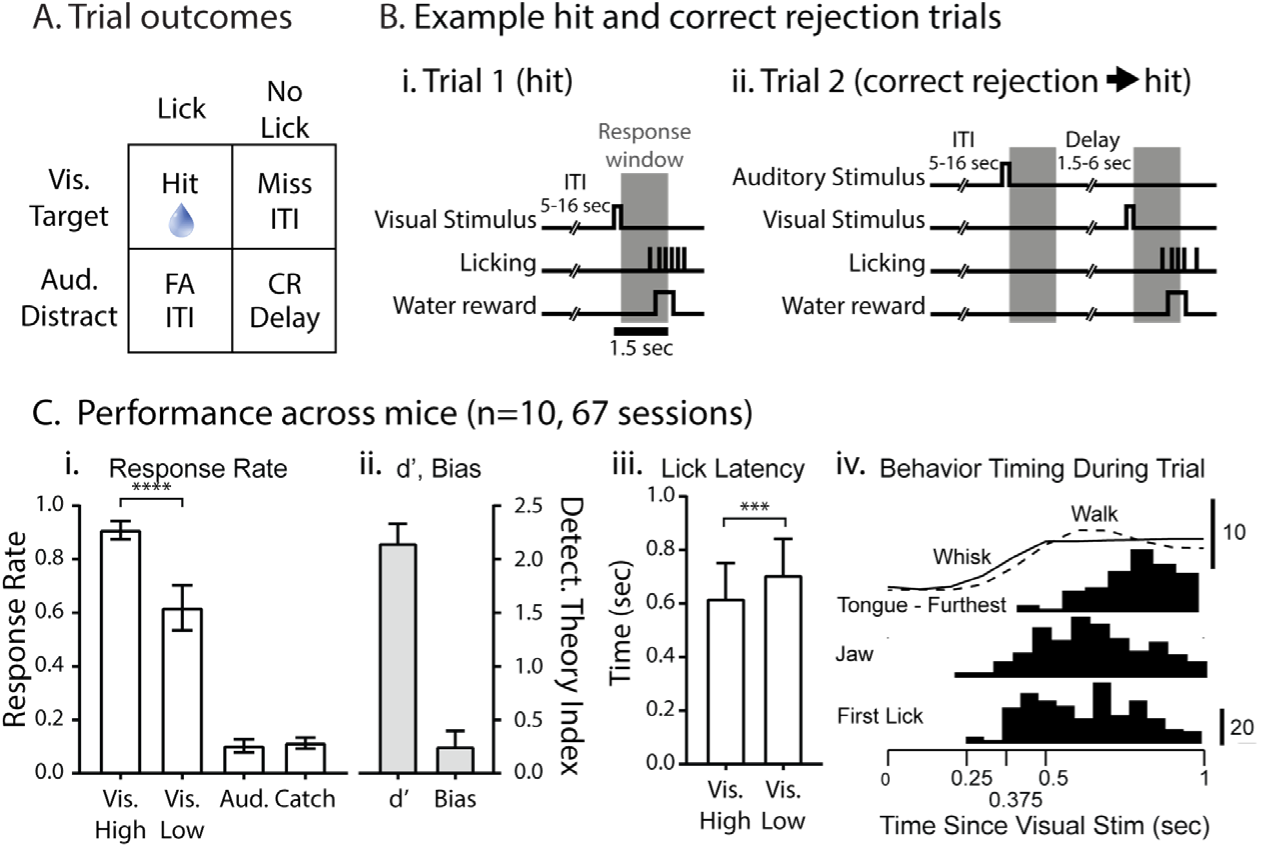
Expert performance and stereotyped behavior during a visual detection task. **A,** Trial outcomes. Either a visual (target) or auditory (distractor) stimulus is presented after an inter-trial-interval (ITI). Mice receive a water reward if they lick within 1.5 seconds of the target (hit). Withholding licking after a distractor (correct rejection) is rewarded with a target trial after a short delay. **B,** Correct examples for target (**i.**) and distractor trials (**ii.**) showing timing of response window and delays. **C,** Response rates (**i.**) and detection theory measures (**ii.**) for all mice (n=10, 67 sessions). **iii.** Lick response times were significantly slower for the low-intensity target stimulus. **iv.** Timing of behavioral responses after visual stimuli. Animals responded initial with increases in whisker movement, and occasionally changes in walking speed (see Fig. S1). Timing of the first detected movements of the jaw, detection of the lick, and the time frame of the furthest movement of the tongue are plotted for 4 mice.

### Training and behavior

Mice were trained in a visual detection task in multiple stages. After learning the task in a freely-moving setup, the behavior was transferred to a head-fixed configuration. The training stages were as follows:

Stage 1: Classical conditioning (1-2 days). An LED flash (target) or an auditory stimulus (distractor) was sporadically presented. The target was paired with water delivery. Mice had the opportunity to drink (9 µl) for 2 seconds before the water was removed from the reward port with suction. The auditory distractor was not paired with water.

Stage 2: Operant Conditioning (1-2 days). Same as stage 1, but water delivery was contingent on licking the reward port within 2 seconds of visual target presentation (response window).

Stage 3: Operant Conditioning with timeouts (3-4 days). Same as stage 2, but licking during the inter-trial-interval (ITI) was discouraged by resetting the time until the next trial. ITIs were randomly chosen from an exponential distribution with min/mean/max of 5/9/16 seconds. Importantly, ITIs were not re-chosen after timeouts, which forced mice to occasionally wait long periods before the next trial.

Stage 4: Full task (6 days, then again after head-post surgery for re-training and imaging). The response window was shortened to 1.5 seconds. Target presentation always occurred after correct rejection of the distractor, but otherwise had a 20% chance of being presented. This reinforced withholding after distractors but motivated mice even when false-alarm rates were high during training. ITIs are same as stage 3 except ITIs after correct rejections were shorter (1.5/2.5/6 min/mean/max exponential distribution). Catch trials were randomly presented instead of stimuli for 15% of trials to assess the spontaneous response rate. For correct example trials see Fig. 1B.

Nine out of 10 mice left training stage 4 with d’ above 0.9. The mouse below this level achieved a high d’ during re-training on the head-fixed configuration. Re-training after head-posting took as few as 2 days. Hit rates were generally higher during the head-fixed version of the task, and this was probably due to the fact that they were less distracted by non-task related behavior, such as exploring the cage.

### Surgery

After training mice were allowed *ad libitum* water overnight. They were anesthetized with isofluorane (0.8-1.5%) and hair on the head, neck, and upper back were removed with a depilatory cream (Nair). Back hair could reflect light on the cranial window during imaging and was removed periodically to prevent noise. Mice were injected with an analgesic (Meloxicam, 1 mg/kg) and antibiotic (Baytril, 5 mg/kg). The skin on the head of the mouse was removed and connective tissue removed until the skull was clean and dry. A custom-made “winged” head-post was glued to the skull with dental cement (3M RelyX Unicem Aplicap). These head-posts had a thin piece of metal on all sides of the transcranial window and were slightly bent to accommodate the curve of the skull. After ∼5 minutes to cure, a small amount of cyanoacrylate (Slo-zap) was placed in the center of the skull which then spread out. Care was taken to avoid bubbles in the cyanoacrylate. A cyanoacrylate accelerator was then applied (Zap Kicker). The imaging window was protected by applying a silicone elastomer (WPI, Kwik-Sil) which was removed each day before imaging and then re-applied. The transcranial window, including skull, became transparent after curing (∼24 hours). Mice were given two days to recover from surgery and water restricted on the third day in preparation for habituation to head-fixation on the treadmill and re-training to the visual detection task.

For combined electrical and optical recordings, mice were anesthetized with isofluorane, ketamine (90 mg/kg, intraperitoneal [i.p.]) and xylazine (10 mg/kg, i.p.). A small (< 0.5 mm in diameter) craniotomy and durotomy were applied in parietal cortex. Post-surgery, mice were transferred for immediate recording.

### Muscimol injection

The unilateral (contralateral to visual stimulus) injection site was chosen by computing a lick rate correlation map of dorsal cortex and targeting the region of high correlation in secondary motor cortex (MOs). Vasculature was used as landmarks. Mice were anesthetized with isoflurane and a dental drill was used to make a small craniotomy at the targeted location in MOs. Pipettes were back-filled with mineral oil and front-loaded with either saline or 2 mM muscimol. The pipette was slowly placed in the cortex 750 µm below the dura and a nano-injector was used to inject approximately 0.4 µl of saline or muscimol over ∼15 minutes. Mice were given ∼2 hours to recover from anesthesia before behavioral testing. This volume of muscimol injection was chosen to inactivate motor cortical regions around the injection site without disruption of activity in nearby cortical areas such as somatosensory cortex (Fig. 4D).

### Electrophysiology

LFP/MUA signals were obtained with 16 channel multielectrode arrays (A1×16-Poly2-5mm-50s-177, NeuroNexus). The probe was positioned at an approximately 45 degree angle to the skull and inserted such that the recording sites were in layer 5 (500-900 µm deep). This positioning enabled the simultaneous recording of layer 5 electrical signals and the overlying through-skull optical signals. Signals were processed through a preamplifier (Multichannel Systems) and amplifier (A-M Systems 3500), bandpass filtered between 0.3 Hz and 5 kHz and digitized at 20 kHz (Power 1401, CED). Spike-field analyses were conducted offline in MATLAB. Electrical signals were high-pass (100 Hz) filtered for spiking analyses and band-pass (20-80 Hz) filtered to isolate the gamma-band LFP signal. Multiunit spike times were identified as threshold crossings well isolated (>2× noise amplitude) from background.

### Imaging

The SNAP25-2A-GCaMP6s mouse line expresses GCaMP6s pan-neuronally, in both excitatory and inhibitory neurons throughout all layers and regions of the cerebral cortex and brain (see Fig. S7E in (Madisen et al. 2015)) and (http://search.brain-map.org/search/index.html?query=Snap25-2A-GCaMP6s). The precise source of widefield signals as measured through the skull are likely to be varied, including potential signals from all layers of the cortex and axonal inputs that arrive from other brain regions, although cortical layers near the skull have a particular advantage owing to proximity to the light source and camera (Mitra et al. 2018; Ma et al. 2016). We do not assign a particular neuronal source to the widefield signals that we measure, but rather assume that they are representative of general neuronal activity within that region of the cortex. We did not observe aberrant (e.g. epileptiform) activity, either optically or electrophysiologically, in our mice, as occurs in some mouse lines expressing GCaMP6 (Steinmetz et al. 2017).

Mice were head-fixed under a macroscope with 75 mm lens (RedShirt Imaging, Macroscope-IIA; 4.5 f; NA 0.4). The cranial window was illuminated with 490 nm light from a mounted LED (Thorlabs M490L4), dispersed with a collimating lens (Thorlabs ACL2520-A). The 490 nm illuminating LED light passed through a 470 nm bandpass filter (ET470/40x, Chroma), was delivered to the skull through a 495 nm long-pass dichroic mirror (T495lpxr, Chroma), and the returning light from the brain (GCaMP6s fluorescence or green reflectance, depending on excitation light) passed through a 510 nm bandpass filter (ET510-32mm, Chroma) prior to entering the camera. In control experiments measuring green reflectance, the cranial window was illuminated with 530 nm LED (Thorlabs LEDC13 530 nm) and a collimating lens, aimed at the surface of the skull. Images were captured with a CCD camera (pco.sensicam QE; 12-bit depth; 1376×1040 pixels). Pixels were binned 8×8 (final resolution of approximately 40-50 um/binned pixel) and images were taken at 20 Hz with Camware software, triggered externally by a MATLAB script running the behavioral protocol. TIF movies were subsequently imported to MATLAB for pre-processing and analysis.

### Fluorescence pre-processing

We desired a readout of neural activity in response to stimuli as well as activity in between trials not directly task-related. A preliminary analysis normalizing raw fluorescence by the mean of each pixel proved unsuitable due to slow changes in the reflectance of the cranial window (e.g. caused by the evaporation of saline on the preparation). We therefore chose to normalize pixel fluorescence by the local mean, computed every two seconds, of ±4 minutes using the calculation ΔF/F=(Fi-F0)/F0 for every pixel where Fi is the raw fluorescences of the ith video frame and F0 is the local mean. This method yielded results similar to calculating ΔF/F for each trial using pretrial activity (Fig. S3). Each movie frame was smoothed with a Gaussian kernel with 3 pixel standard deviation (120 μm). We call this readout ΔF/F throughout the paper. See https://github.com/salkoffd/PreProcessWfRecording for MATLAB code.

### Brain atlas alignment

We manually aligned one calcium recording (average picture) per mouse to the Allen Mouse Common Coordinate Framework (Oh et al. 2014) (Fig. S2) using landmarks such as bregma and the divergence of the cortical hemispheres at the anterior aspect of the superior colliculus. Subsequent recordings were automatically aligned to the reference image for that mouse with image registration in MATLAB.

### Behavioral analysis

The absence of licking on a Go trial could indicate a miss, or alternatively the mouse is disengaged from the task. We therefore only analyzed miss trials during “motivated” periods, which were defined as epochs of licking no further than 70 seconds apart. Motivated epochs of less than three minutes duration were omitted.

Response (hit) rate is the fraction of visual stimulation trials in which the animal successfully licked during the response period (Fig. 1). Discriminability (d’) and bias were calculated as previously described for signal detection theory (Swets and Green 1978). Discriminability (d’) was calculated as norminv(hit_rate)-norminv(FA_rate) where norminv is the inverse of the normal cumulative distribution function. Bias (criterion) is computed as - 0.5*(norminv(hit_rate)+norminv(FA_rate)). We refer to task performance as a constellation of these behavioral measures (hit rate, false alarm rate, d’ and criterion), as per signal detection theory.

Video of the face and pupil was acquired at 10 Hz with a Basler acA1300-30um. Pupil position and diameter was analyzed with a supervised script (McGinley, David, and McCormick 2015). Eye movement was quantified by finding the 2-point-smoothed pupil position before and after each frame and calculating the hypotenuse.

Running on a treadmill was measured at 10 Hz using an optical mouse. Spontaneous run bout initiations were defined as moments when the treadmill velocity exceeded 2 cm/s for ≥1 second immediately after the absolute velocity was below 2 cm/s for ≥1 second. Spontaneous run bout cessations were found by flipping the treadmill velocity vector and using the same method.

Hit trials were subcategorized into those associated with increased running, decreased running, or neither. First a minimum and maximum treadmill velocity were found in the first 0.5 seconds after the target stimulus. Trials were labeled as increased running when the max occurred after the min and was ≥5cm/s larger. Trials were labeled as decreased running when the max occurred before the min and was ≥5 cm/s larger (Fig. S1, S8F).

### Activity difference maps

We computed the difference in activity (ΔF/F) between post and pre-stimulus time-points, (Fig. 2B), between different light conditions (Fig. S6B), and between trial types (Fig. 3A). Calculating the significance for each pixel in a large image poses a multiple comparisons problem. We therefore used a permutation test (Nichols and Holmes 2002) to calculate the significance per pixel as well as significance for the image (evaluating both the total number of pixels with p<0.05 and the size of the largest cluster of pixels with p<0.05). This allows for an estimation of image statistics under the null hypothesis (trial types are not different) and does not make assumptions about the shape of the data. We randomly permuted trial labels 1000 times, each time computing the difference between trial types, and defined significant pixels as in the ≥97.5^th^ or ≤2.5^th^ percentile of the permuted data. The permuted datasets were used to acquire distributions for the total number of significant pixels and size of the largest cluster under the null hypothesis, to which the actual dataset was compared. To avoid counting clusters containing pixels with both negative and positive values, we searched separately for significantly positive and negative clusters. See https://github.com/salkoffd/ActivityDifferenceMap for MATLAB code. Due to high computational expense, we first downsized images to 25%, then upsized difference maps before projecting them onto the common coordinate framework. We analyzed the activity difference only within the cortical region of interest (ROI, excluded midline, ipsilateral cortex, and regions outside the cranial window), however for illustrative purposes we included the full imaging window in some figures (e.g. Fig. 2B, 4D).

**Figure 2.**
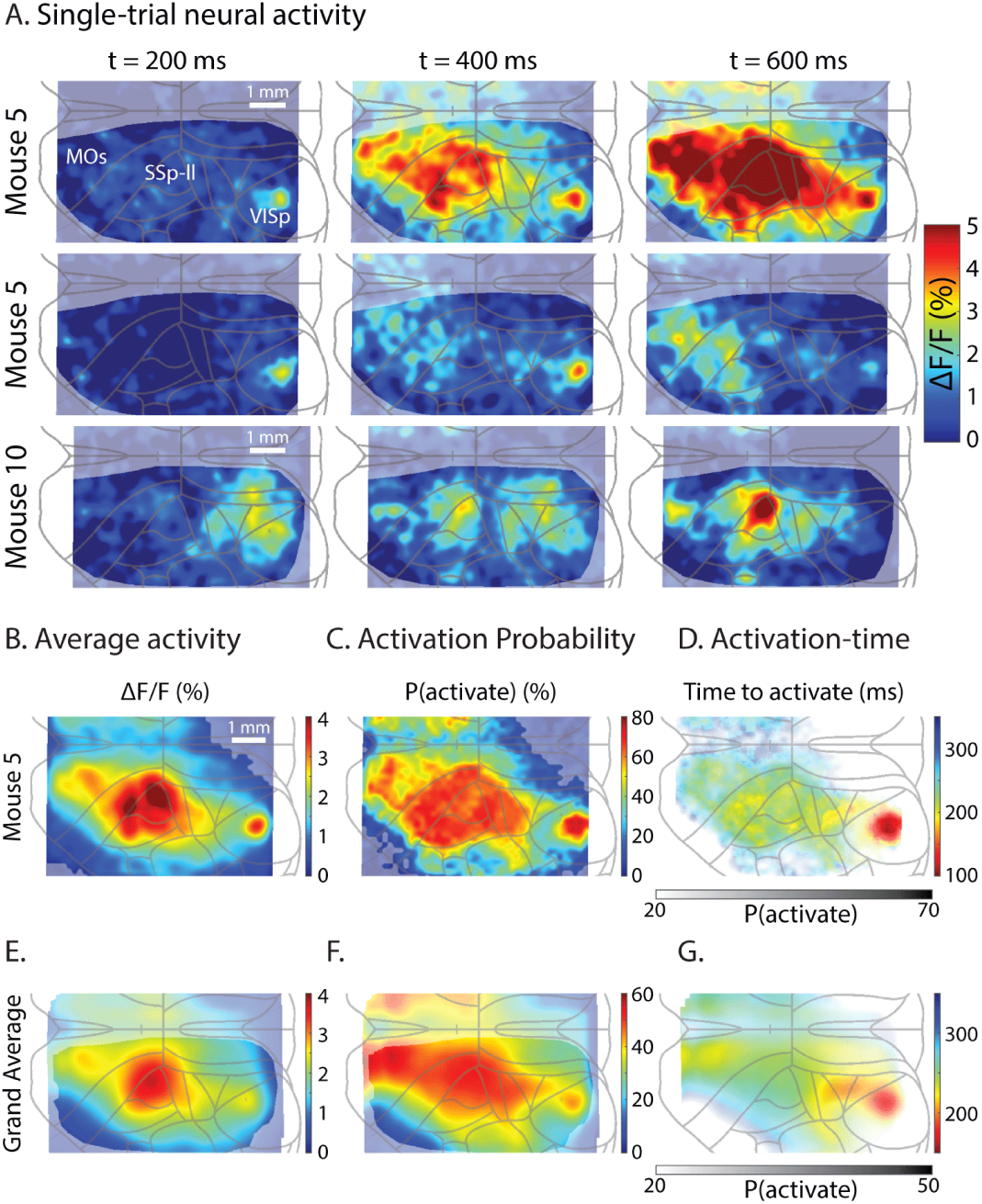
Widespread task-related neural activity. **A**, Single-trial neural activity on hit trials after target presentation (t = 0 ms). Each row is a separate trial and two trials are shown for mouse 5. The response times for the three example trials were within 10 ms of each other (mean 0.524 seconds). Activity was highly variable in time and space. Scale bar for mouse 5 and 10 shown. **B**, Average activity (ΔF/F) after t = 450 ms for an example session. Trials with response time faster than 500 ms were omitted from the analysis. Opaque pixels show a significantly large cluster of pixels with p<0.05 (p<0.001, permutation test between pre and post-stimulus activity). **C**, Activation probability after t = 450 ms, defined as the probability each pixel exceeds threshold (derived from pre-stimulus values) per trial, (see methods). Opaque pixels show a significantly large cluster of pixels with p<0.05 (p<0.001, shuffle test). Same trials as B. **D**, Activation-time, defined as the average time to exceed a low threshold. Note activity first arises in VISp. Opacity is derived from C. Same session as B and C. **E-G**, Same as B-D but averaged across all mice. Opaque region of E and F show cortical ROI common to at least 5/10 mice. Scale bar in E is averaged since images were stretched to match a common coordinate system (see methods). We adopt here the abbreviations of the Allen Institute Common Coordinate Framework (Fig. S2; (Oh et al. 2014)). MOs - secondary motor cortex; SSp-ll - primary somatosensory cortex, lower limb; Visp - primary visual cortex.

**Figure 3.**
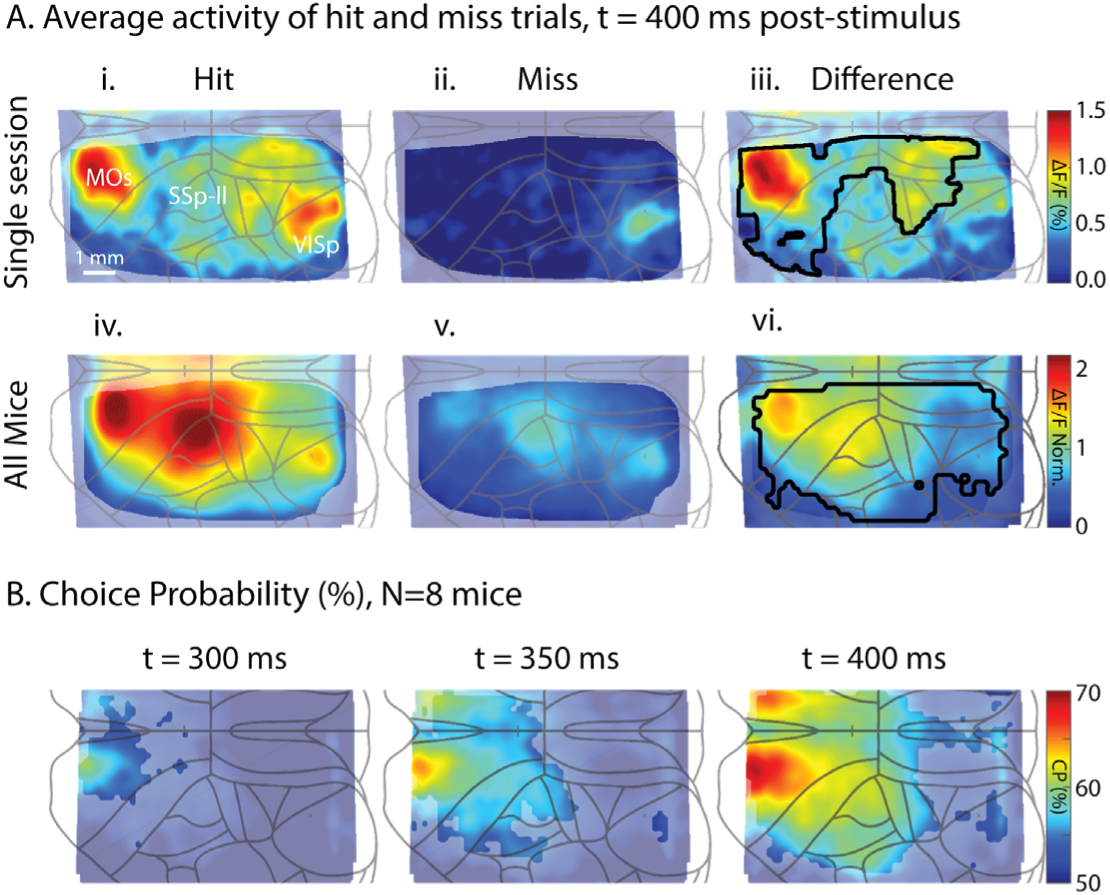
Choice-related activity arises in secondary motor cortex. **A**, Average activity of hit and miss trials 400 ms after low-intensity stimulus onset. hit trials with response time faster than 450 ms were omitted from the analysis. **i-iii** is from one session (n=26 hit, 20 miss trials), **iv-vi** is the average of 8 mice. Black outlines in difference maps are clusters of pixels (each pixel p<0.05) significantly larger than that generated by permuted maps (p<0.001, permutation test). In iv-vi, ΔF/F has been normalized to account for differences in fluorescence across mice, but preserves the difference between hit and miss activity within each mouse. **B**, Choice probability (CP) calculated across time. Opaque pixels have CP significantly larger than chance (>50%, p<0.05, one-tailed t-test). Note origin of CP in MOs and subsequent caudal spread. Only trials with response times >450 ms were analyzed. Median response time 732 ±120 ms across mice.

For comparing hit and miss activity across mice (Fig. 3A), we first normalized ΔF/F. This was done because overall intensity changes varied between mice. The normalization factor was computed by first finding the cortical region imaged in all 10 mice. Then, for each mouse, we found the average pixel value within that region including both hit and miss activity. Figure 3A is therefore in units of the average cortical pixel within mouse (0.62% ± 0.49 ΔF/F).

### Choice probability maps

Choice probability is a signal detection theory measurement describing how well an ideal observer can predict the decision of an animal (Britten et al. 1996). It is calculated by finding the area under the ROC (receiver operating characteristic) curve, and in the context of this study is the probability that the activity on a hit trial is larger than on a miss trial, if one of each trial type is selected randomly. Similar to the activity difference map, we used a permutation test to calculate significance for each pixel and for the image (Nichols and Holmes 2002). Trial labels were permuted 1000 times, each time calculating the choice probability. The actual choice probability was then compared to this distribution to derive a p-value. The permuted datasets were used to acquire distributions for the total number of significant pixels and size of the largest cluster under the null hypothesis, to which the actual dataset was compared. See https://github.com/salkoffd/ChoiceProbabilityMap for MATLAB code. Due to high computational expense, we first downsized images to 25%, then upsized choice probability maps before projecting them onto the common coordinate framework.

### Activation probability maps

Activation probability provides a measure of how much more likely neuronal activity increases after a stimulus compared to the chance of increasing in the absence of a stimulus. For each pixel, we calculated the probability of activation above chance:

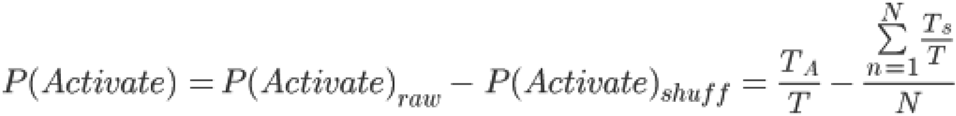

where T is the number of trials, T_A_ is the number of trials on which the pixel became active, N is the number of iterations (1000), and T_S_ is the number of shuffled trials on which the pixel became active. A pixel is “activated” for the trial if it exceeds a threshold of 3.96 standard deviations above the mean of pre-stimulus values (1 sec, 20 frames) within 500 ms of stimulus onset. This threshold corresponds to *α* = 0.05 with a Sidak correction for multiple comparisons: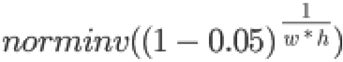 where w * h is the size of the (downsized) video (32*43).

The “raw” activation probability 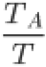, was corrected since each pixel has a baseline chance of exceeding threshold (due to spontaneous activity, arousal, etc.). We therefore shuffled the time of analysis (±2 minutes maximum relative to each trial) N=1000 times. The shuffled data was used to find the statistical significance of each pixel. Shuffled data also provided distributions of pixel cluster sizes under the null hypothesis (stimulus has no effect), to which the actual data was compared. See https://github.com/salkoffd/pActivateMap for MATLAB code. Due to high computational expense, we first downsized images to 25%, then upsized activation probability maps before projecting them onto the common coordinate framework. We calculated activation probability only within the cortical ROI, however we used the entire imaging window in Fig. 2C-G for illustrative purposes.

### Activation-time maps

The activation-time map reveals the time course of initial activity rises after the stimulus. It is essentially the average time required to reach a low threshold for each pixel. For each trial and each pixel, we first calculated the mean and standard deviation of pre-stimulus values (1 sec, 20 frames). Pixels were further considered if they crossed a high threshold (>4.58 standard deviations above the mean, derived from setting alpha=0.05 and applying the Sidak correction for the number of pixels in each frame of the movie, 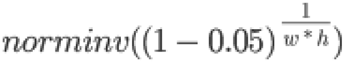 (where w*h is 130*172) within 1.9 seconds post-stimulus. If this high threshold was crossed we found when the lower threshold was crossed for two consecutive frames (85^th^ percentile of pre-stimulus values). These times were then averaged across trials.

### Correlation maps

Lick correlation maps (Fig. 4C) were generated by computing lick rate in 0.1 second bins, then smoothing by ±3 points. Run speed correlation maps (Fig. S8D) were generated by smoothing running by ±4 points. Pupil diameter correlation maps (Fig. S8E) were generated by smoothing pupil diameter by ±3 points. The maximum correlation for each pixel was found within ±700 ms.

**Figure 4.**
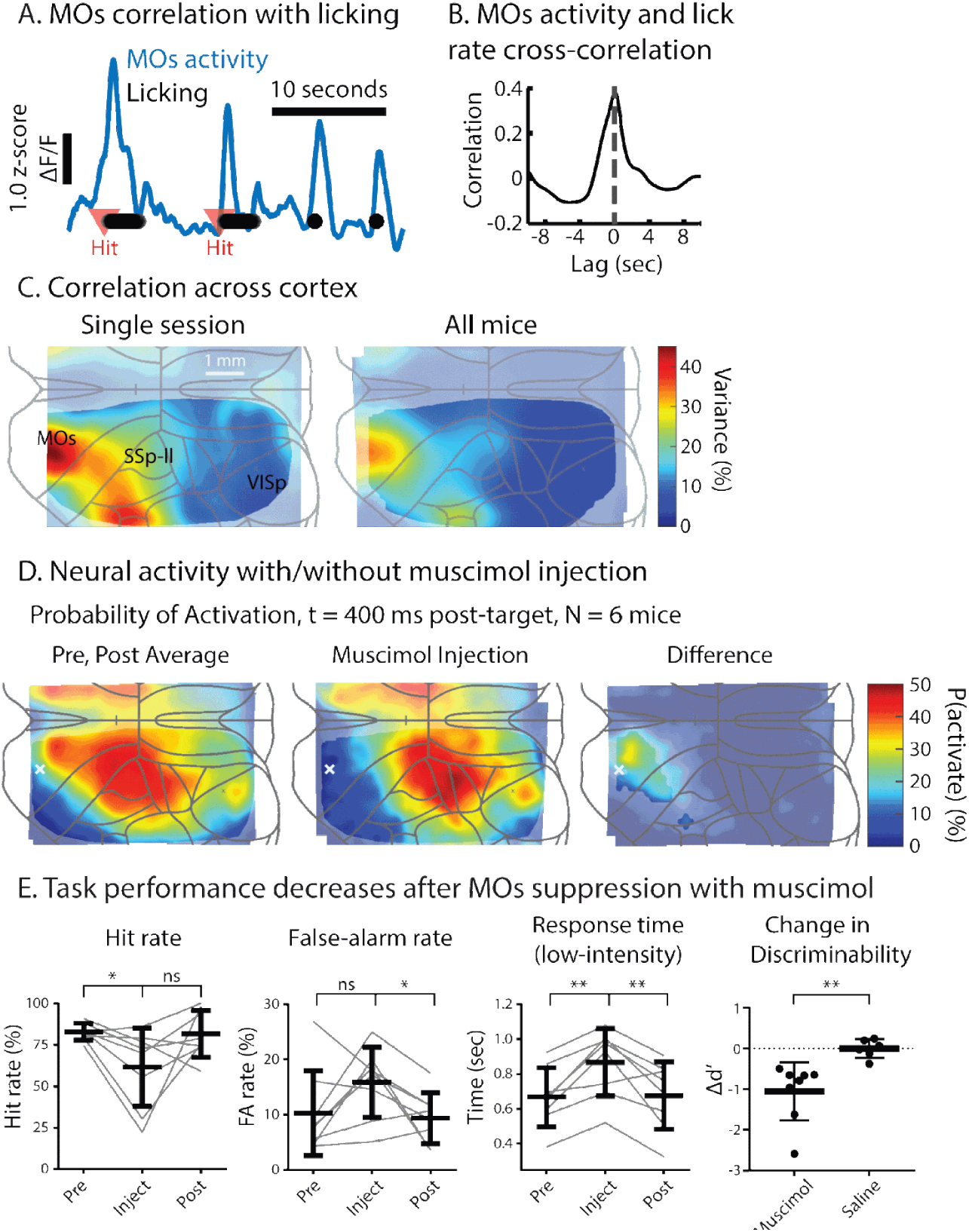
Secondary motor cortex (MOs) is critical for goal-oriented licking. **A**, Example of MOs activity alongside lick times (black dots). **B**, Cross-correlation of MOs activity and lick rate for an example session. **C**, Neural variance explained by lick rate across cortex for a single session (left) and averaged across all mice (right, n=10, 67 sessions). **D**, Neural activity with/without muscimol injection. Probability of activation 400 ms after target onset for 6 imaged mice. White x is the approximate location of the injection. Opaque pixels in the difference map are significantly greater than 0 (p<0.05, two-tailed test). **E**, Task performance decreases after MOs suppression with muscimol (n=8 mice). Muscimol-injected mice have a reduction in hit rate and an increase in false alarm rate, resulting in a significant reduction of discriminability (d’) compared with saline-injected mice.

### General linear model predicting response on target trials

The logistic binomial model (Fig. 5) included all (motivated) target trials with response time larger than 450 ms. We included six predictors: high versus low-intensity target, MOs post-stimulus response (350-450 ms), pre-stimulus eye movement (−300 to 0 ms), pre-stimulus pupil diameter (−300 to 0 ms), run speed at target presentation, and average pre-stimulus activity of the entire cortical ROI. Sessions were combined per mouse to maximize the number of data points in each GLM. All predictors were converted to z-score before fitting. The data was fit with custom MATLAB script and the fitglm function with the linear model specification and binomial distribution for the response variable. The full models were compared against models generated after permuting trial labels (i.e. hit/miss, 1000 iterations). For reduced models, we permuted predictors instead of removing them to control for number of predictors (1000 iterations). Model fit was assessed with d’ and accuracy (defined as the average of hit and miss correct classification probability).

**Figure 5.**
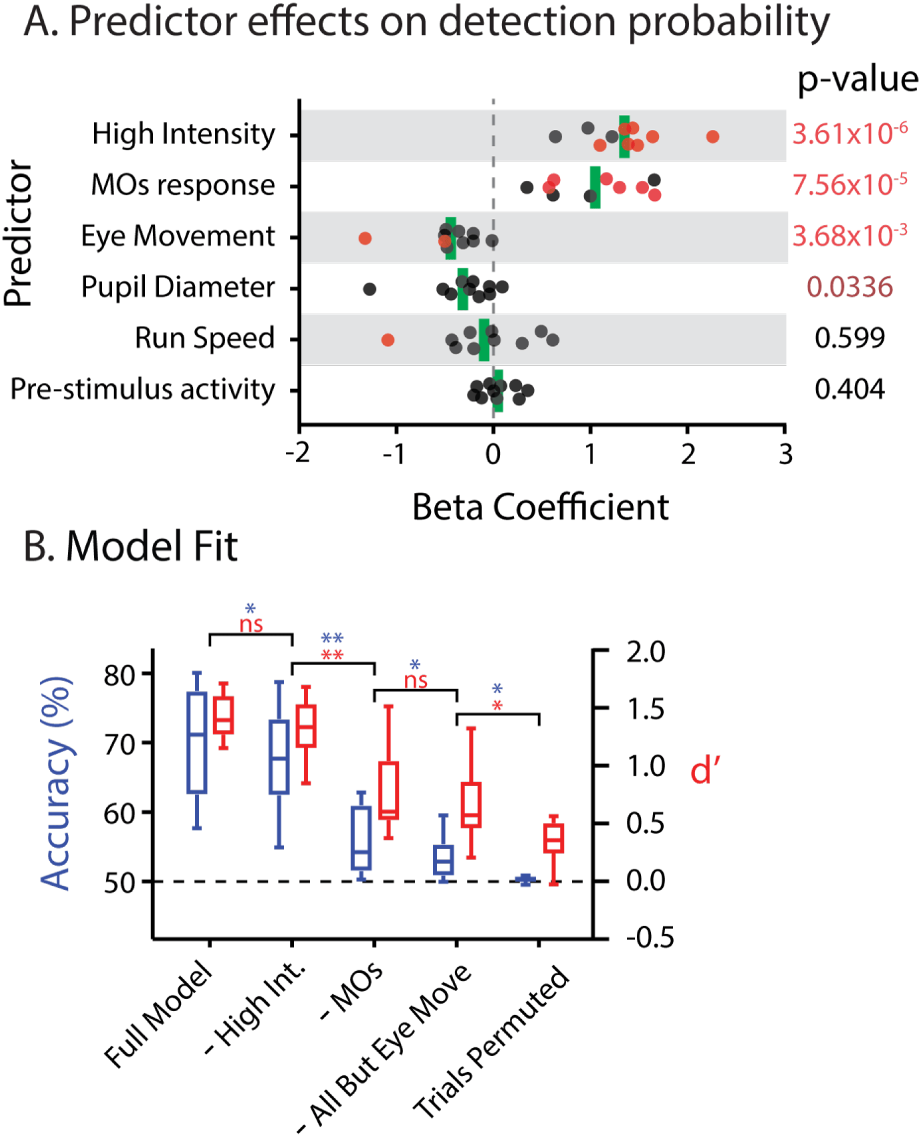
Secondary motor cortex (MOs) is a robust predictor of target detection. **A**, Logistic regression was used to find the effects of several predictors on the probability of licking. Trials with response times <450 ms were omitted. While the high-intensity stimulus and MOs activity (400 ms post-stimulus) increased the likelihood of licking, eye movement (≤300 ms before the target) and arousal (pupil diameter at target onset) decreased the likelihood. Run speed and pre-stimulus cortical activity had no effect. Red points indicate the predictor was significant within mouse after the Sidak correction for multiple comparisons. **B**, Model fit was assessed with prediction accuracy and d’ (see methods). The full model explained behavior well. Elimination of predictors via permutation reduced performance. **–** High Int.: high/low intensity predictor was permuted. **–**MOs response: the MOs post-stimulus activity was also permuted. Eye move only: All predictors except eye movement were permuted.

### Linear model predicting neural response during trial behavior

For each session (n=37 from 6 mice), a design matrix was constructed by extracting 10 features for each trial (Fig. 6). These features included values representing details of the trial structure, behavioral response, and facial movements extracted from the video (Fig. 6A,B). Trials with licking before 400 ms were omitted from the analysis. A feature vector was created representing each 0.1 s for the 2 s after each stimulus was presented. The features extracted from the video include video motion energy (ME), calculated as the sum total of absolute change in pixel intensity between adjacent video frames, whisk ME, calculated in the same manner as video motion energy, except for a region restricted to the main whisker pad (Fig. 6B), pupil diameter, and eye movement (calculated as indicated above). The analog features from the video and the wheel velocity, were smoothed by taking the average of the current data point and 0.3 seconds prior. The trial variables, hit, miss, false alarm, response time of first lick, light presentation intensity (higher or lower) and stimulus type (auditory, visual or no stimulus) were constant throughout each trial. Time was relative to when the last stimulus was presented.

**Figure 6.**
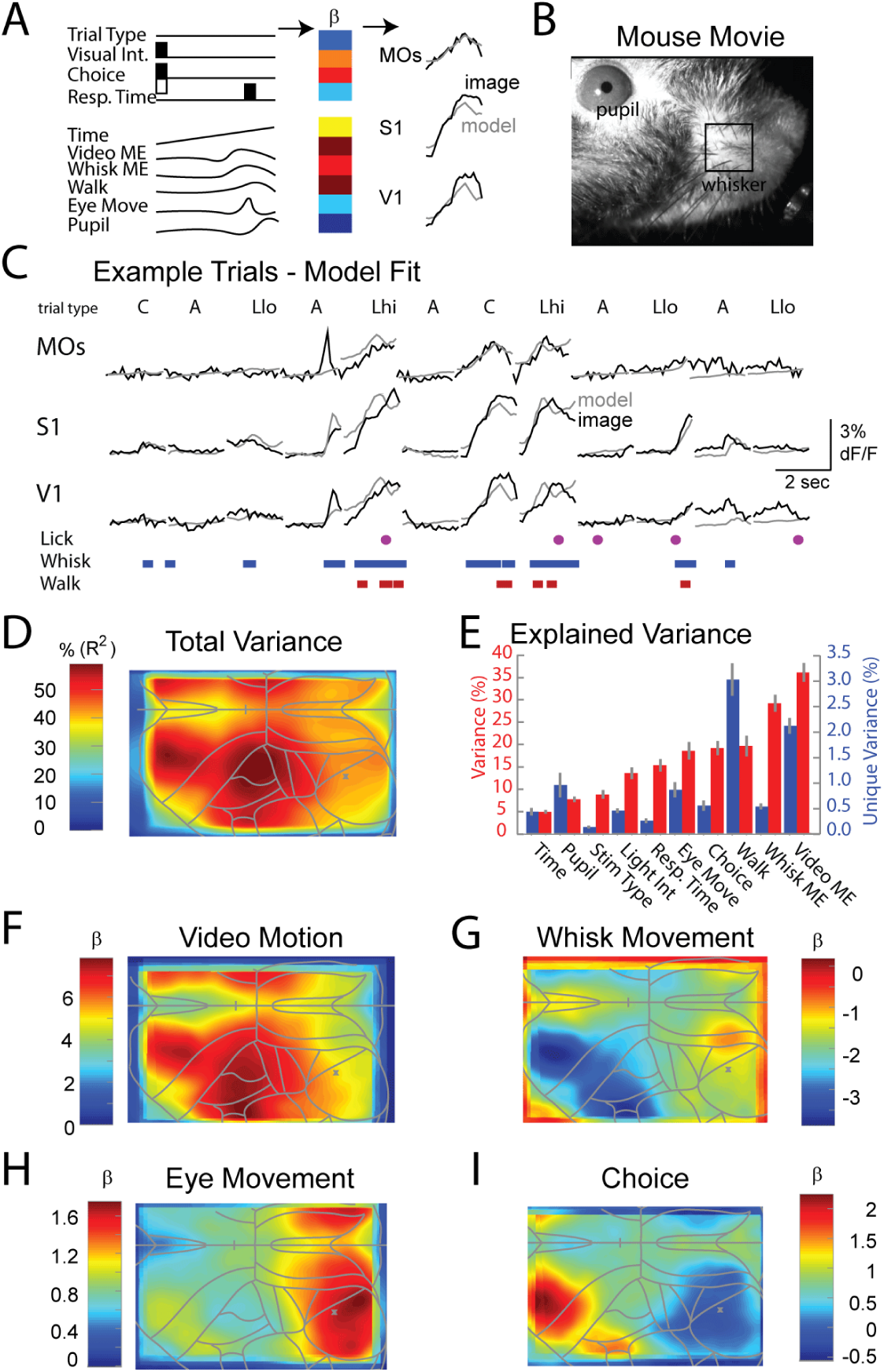
Widespread cortical activation is largely explained by movement and response choice. A, a ridge regression linear model (see Methods) was used to fit the pixel-by-pixel amplitude-time course for each trial (37 sessions, 6 mice). Regressors used in the model include unitary variables: trial type (visual, auditory, catch), intensity of visual stimulus (low or high), response choice (hit/no response/false alarm), response time and continuous variables: time, motion energy in the entire video, motion energy in the whisker pad only (see B), walking speed, eye movement amplitude, and pupil diameter. The linear model generated spatiotemporal maps of β weights for each variable (see D-G) and were used to predict the amplitude-time course of each pixel of the widefield movie of cortical activity. Examples of the activity and model fit for averaged regions of pixels in MOs, S1, and V1 are shown (see C). B. Example frame from a video of the mouse face and eye during the performance of the task. The video was used to examine movements of the face (video ME), whisker pad (whisk ME), eye, and pupil diameter. C. Example fluorescent traces and the model fit for regions of cortex in MOs, S1, and V1. Trial type was C=catch (no stimuli), A=auditory, Llo=lower intensity light; Lhi=higher intensity light. Periods whisking and walking, and time of first lick, are illustrated. Note that whisking is associated with widespread cortical activity that is well fit by the model. D. The model is able to explain a significant fraction (approximately 35-55%) of variance of neural activity during performance of the trial. E. Total and unique explained variance for each parameter. Movement (video, whisk, walk, eye movement) explains a high degree of neural activity, while other variables such as arousal (pupil diameter), response choice and timing, also make significant contributions. Bars are mean+/-SEM. F-I. Spatial maps of the β weights of the model for video motion, whisk movement, eye movement, and behavioral choice. Note that behavioral choice peaks in MOs. The model was run on data from 6 mice in which the video contained images from the nose to just behind the eye and from above the eye dorsally to the lower lip ventrally.

### Ridge regression model

To reduce the impact of the multicollinearity of these features, we used Ridge Regression, instead of the typical general linear model, and the optimal penalty term was found through cross validation. To account for overfitting, the model was fit and evaluated for explained variance (R^2) on cortical pixel activity for each session using 10-fold cross-validation. To compute the unique contribution of each feature, we permuted one feature at a time to create a reduced model. The unique explained variance was the difference between the full model and the explained variance of the reduced model. To find the unique variance of multiple features combined, a similar method was used but multiple features (e.g. movement) were permuted together.

### Statistics

Statistical tests were performed in either MATLAB 2015b or GraphPad Prism 7.01. Reported p-values are calculated from permutation tests unless otherwise stated. Figures use following notation: * p<0.05; ** p<0.01; *** p<0.001; **** p<0.0001.

## Results

### Expert performance and stereotyped behavior during a visual detection task

In order to understand the encoding of behavioral and task variables throughout cerebral cortex, we trained mice (n=10) in a simple Go/No-Go visual detection task (Fig. 1A, B). Mice were required to lick a reward port after presentation of either a dim or less dim, 50 ms LED flash (target) and withhold licking after an auditory sweep (distractor, Fig. 1B, see methods for details). Licking after target presentation (hit) was rewarded with water and withholding after the distractor (correct rejection) was rewarded with a short ITI and subsequent target trial (Fig. 1B). After training in a freely-moving setup and habituation to head-fixation on a treadmill, mice performed expertly (Fig. 1C), with 90.9% (±3.42) hit rate for the high-intensity target and 10.3% (± 2.47) false-alarm rate, yielding an overall discriminability (d’) of 2.18 (±0.21).

Awake mice exhibit constant fluctuations in arousal and movement (McGinley et al. 2015; Musall et al. 2019; Stringer et al. 2019), however the relationships between these fluctuations and task performance are poorly understood. We measured treadmill speed, pupil diameter, whisker pad and jaw movement, and furthest tongue extension, while mice performed the visual detection task. We found that following presentation of the visual stimulus target, animals often respond first with movements of their whiskers, and sometimes with either increases or decreases in walking speed (Fig. 1Civ and Fig. S1). These changes in whisker/walk movements were followed by opening of the lower jaw and detection of a lick over the next 1-2 seconds with the earliest consistent first lick latency of approximately 375 msec (Fig. 1Civ).

### Widespread, task-related neural activity

Which cortical regions are active following target presentation and during response initiation and performance? We used wide-field GCaMP6s imaging of pan-neuronal intracellular Ca^2+^ to monitor activity in the dorsal aspect of the left (contralateral to visual stimulus) cortex as mice performed the task. The imaging window spanned from primary visual cortex (VISp) caudally to secondary motor cortex (MOs) rostrally, allowing for observation of a wide proportion of occipital, parietal, and frontal cortices (see Fig. S2 for details on image alignment and region acronyms). Following presentation of the light, increases in neural activity appeared first within visual cortex (Fig. 2). Over the next several hundred milliseconds activity spread to include somatosensory and motor cortical regions, corresponding to the timing of the changes in walking, whisking, and other orofacial movements associated with performance of the task (Figs. 1C, 2A; see below). Comparison of single-trial cortical activity revealed a variety of spatial patterns shortly after target presentation. However, hit-related activity averaged within a session revealed consistent activation of frontal and parietal cortices (see Fig. S4 for examples from 6 additional mice; see supplemental movie). To reveal this spatial activation pattern, we computed an “activation probability map”, which is the probability (above chance) that any one region becomes active on a hit trial (≤450 ms post-stimulus, see methods) (Fig. 2C for a representative session and Fig. 2F average of n=10 mice and n=67 sessions). We defined activation per pixel as activity significantly above a trial-by-trial threshold set by the statistics of pre-stimulus activity for that pixel (see methods). 94% of sessions were significant for the number of active pixels in the imaged region of cortex (p<0.05) and 91% of sessions had clusters of high activation probability significantly larger than expected by chance (p<0.05), indicating that widespread cortical activation was typical for target detection. We observed a nucleus of high activation probability in VISp corresponding to the approximate expected retinotopic location of the visual stimulus (Fig. 2B-G). High activation probability extended from visual areas to secondary motor cortex and included much of somatosensory cortex. Lateral aspects of the cerebral cortex, including auditory cortex, were outside of our field of view (see Fig. S2).

We analyzed the time course of activation by generating an “activation-time map”. This is the average time activated pixels required to reach a threshold (85^th^ percentile of pre-stimulus values, see methods). A representative session (Fig. 2D) and grand average (Fig. 2G) are shown. As expected, VISp activated first after target presentation (145 ms ± 32.5; calculated for the VISp activity hot-spot on a per mouse basis). Some mice had activation-time maps with gradients pointing towards MOs (Fig. 2D) while in other mice activity seemed to jump quickly to retrosplenial cortex and MOs (evident in Fig. 2G). VISp became active significantly earlier than MOs (78.7 ms, p=2.83×10^-4^, paired t-test of hot-spot in VISp and MOs). These findings support widespread dorsal cortical activation between the stimulus and response for a visual detection task.

We performed multiple control experiments to ensure that widespread fluorescence changes were caused by GCaMP6s-mediated fluorescence rather than possible artifacts including spatial scattering of light through the skull (Fig. S5), movement artifact, or otherwise non-GCaMP6s related fluorescence changes (Fig. S6). Our results suggest that, while non-negligible, non-GCaMP6s related signals cannot account for the magnitude and spatial extent of fluorescence increases during target detection.

### Choice-related activity arises in secondary motor cortex

Neural activity during target detection represents a mixture of stimulus and choice/response-related signals. To isolate signals related to choice/response we analyzed the difference between hit and miss trial activity. We analyzed only low-intensity target trials, which had a lower hit rate and slower response times. To reduce contamination of neural signals related to licking and somatosensory feedback of the tongue on the reward port, we analyzed activity for the 400 ms after target presentation and discarded trials where the mouse licked sooner than 450 ms. Widespread activity was present on low-intensity hit trials (Fig. 3Ai example session, Fig. 3Aiv grand average of 8 mice), and was particularly high in MOs and medial somatosensory and motor cortical regions. Miss trials exhibited low level, but significant, activity throughout the cortex, again centered in the visual, somatosensory and motor cortical regions (Fig. 3Aii, v). To quantify the difference between hit and miss activity and control for multiple comparisons we performed a permutation test within each session. Of the recording sessions with at least 10 hit and miss trials, 46% (13/28 sessions) had a significantly large cluster of pixels (each pixel p<0.05) greater on hit than miss trials (example shown in Fig. 3Aiii). MOs was always overlapped by the significant cluster in these sessions. When we looked across all mice we found activity was significantly larger in hit trials than miss trials in frontal, parietal, and much of occipital cortex, and this cluster of pixels (each pixel p<0.05) was significantly larger than expected (p<0.001, permutation test, n=8 mice, Fig. 3Avi). The grand map of the difference between hit and miss trial activity (Fig. 3Avi) shows an increasing difference in the caudal-rostral direction, with the largest difference within a hot-spot in secondary motor cortex. Spontaneous licking during the inter-trial interval was associated with similar spatial patterns of activation, yet of smaller amplitude (Fig. S7), further supporting the role of these regions in behavioral and neural components of response initiation.

Activity in the somatosensory cortex corresponding to lower limb areas (SSp-ll) were found to relate to changes in running speed (Fig. S8). Furthermore, SSp-ll activity increased with reward-related licking (Fig. S7, S8), spontaneous licking (Fig. S7), and orofacial movements (Fig. 6F). At this point it remains unknown to what extent this region is causal for these behaviors.

Examining the differences between miss and hit trials in pre-stimulus cortical activity revealed that miss trials are associated with pre-trial activation of somatosensory and visual cortical regions (Fig. S9A). Presumably, these pre-trial activations correspond to spontaneous movements of the face or eyes, since similar patterns of activity are associated with these movements (e.g. Fig. S9B; n=9). Since our typical measure of cortical response subtracts pre-trial activity, we wondered if post-stimulus trial activity alone varied on hit and miss trials. Examining the hit minus miss post-visual stimulus trail activity alone (without subtraction of pre-trial activity) at 400 msec revealed large activation in area MOs and a decrease in activity in visual cortex, (Fig. S10; n=8). These results suggest that changes in both pre-trial cortical activity as well as post-stimulus cortical activation significantly affect the ability of the animal to perform the task.

To further quantify response-related activity we calculated choice probability (CP) for each recorded pixel. CP is the probability that the activity on a hit trial is larger than on a miss trial if one of each trial type is selected randomly (Britten et al. 1996)(see methods). 100% CP corresponds to perfectly encoding hit trials whereas 50% is chance. Again, we omitted hit trials with response times sooner than 450 ms. At 400 ms post-stimulus we observed high (∼70%) average CP within MOs (Fig. 3B, right panel), with some sessions having >80% CP (not shown). Similar to the hit-miss activity difference map, CP percentage increased along the caudal-rostral axis. We wondered whether this reflected an accumulation of target evidence pooled by MOs or, alternatively, a dispersion of choice signal originating from MOs. We calculated CP for every video frame after the target onset and found significant CP first in MOs (300 ms post-target), followed by subsequent spread to more caudal regions of cortex (Fig. 3B, opaque pixels have p<0.05). These data suggest activity within MOs is first to represent the response decision and that this activity then spreads caudally as the animal’s response is further implemented. Combined, we find a caudal-rostral spread of stimulus encoding (Fig. 2D,G) and a rostral-caudal spread of response initiation and implementation (Fig. 3B).

### MOs is critical for goal-oriented licking

Previous research identified MOs as critical for controlling lick responses during detection tasks (Guo et al. 2014; Zagha, Ge, and McCormick 2015; Goard et al. 2016; Allen et al. 2017; Chen et al. 2017). We wondered if this was the case in our task. We first asked which regions are correlated with licking by calculating the pixel-wise correlation of ΔF/F and lick rate across dorsal cortex. We found two regions of interest: anterior MOs and a somatosensory region spanning representations for the nose and mouth (Fig. 4C; see supplemental movie). This latter region was not particularly active before the lick response (Fig. 3), and we hypothesize that this region becomes active due to sensory feedback from mouth movements and or interaction with the reward port during lick bouts. When activity from anterior MOs is plotted with licking, a correlation is evident for both target-related and spontaneous licking (Figs. 4A, S7). Cross-correlation analysis revealed the width of this correlation to be ∼3 sec (Fig. 4B) which corresponds roughly to lick bout duration. We also observed a high correlation between the lick response time and time for MOs activity to reach an arbitrary threshold (Fig. S11). Out of all imaged cortical regions, response time correlation was highest in MOs. The MOs region was also active during reward-related licking in naive mice, although in naive mice MOs activation occurred 100-200 ms after licking initiation (n=3 mice, data not shown).

Secondary motor cortical (MOs) activity may passively reflect the decision to lick or, alternatively, may be necessary for modulating task-related licking. In order to distinguish between these scenarios we performed unilateral muscimol or saline injections in MOs contralateral to the visual stimulus (Fig. 4D,E). Muscimol is a GABA_A_ agonist and silences neuronal activity. The precise injection site was determined by calculating a lick correlation map for the previous day’s session (example in Fig. 4C) and targeting the region of high correlation within anterior MOs [2.10 +/- 0.29 mm anterior, 1.65 +/- 0.16 mm lateral, similar coordinates to area ALM identified by (Komiyama et al. 2010; Guo et al. 2014) as important for licking]. Mouse performance was tested ∼2 hours after injection to ensure anesthesia had worn off.

To assess the spatial extent of muscimol suppression, we imaged GCaMP6s activity before, during, and after unilateral muscimol injection for all hit trials. We calculated probability activation maps to quantify the degree of suppression (Fig. 4D). As expected, activity at the unilateral MOs injection site during hit trials was strongly suppressed (to ∼5%), and suppression was mostly confined to motor cortex on the side of the injection (Fig. 4D right panel, opaque pixels have p<0.05). Although we did not image the entire contralateral motor cortex during our task, we did observe that at least the medial portion of the secondary motor cortex within the imaging region was active on hit trials, in a manner that was similar to prior or after recovery from muscimol injection (Fig. 4D). Analysis of task performance of muscimol-injected mice revealed a significant decrease in hit rate and increase in false-alarm rate, contributing to a significant reduction in discriminability (d’) compared to saline-injected controls (d’ difference −1.05 ±0.71; p=0.0048, n=8 mice; two-sampled t-test; Fig. 4E, results for saline-injected mice (n=6) shown in Fig. S12). We also found the response time for the low-intensity target was significantly increased by muscimol injection (Fig. 4E, p=0.003, paired t-test of with/without muscimol). The ability of the animal to partially perform the task (at a reduced d’) may be explained by the ability of the contralateral MOs to remain active (Fig. 4D), since each hemisphere may compensate for the loss of the other (Guo et al. 2014; Li et al. 2015). In summary, the high correlation of activity within MOs with licking, the ability of activity in MOs to predict response choice, and the significant decrease in task performance with unilateral MOs inactivation all support MOs as critical for goal-oriented licking.

### MOs activity is a robust predictor of target detection

Activity in MOs was found to be correlated with choice, however it may also covary with other predictors of the mouse’s decision (such as run speed). To test the ability of MOs to predict target detection while controlling for other variables, we performed logistic regression. While considering which predictors to include in the model we found that miss trials were associated with higher pre-stimulus activity within the imaged cortex, owing to either spontaneous cortical rhythms, arousal, or eye movement-related activity (Fig. S9). We therefore included six variables to predict whether the mouse licked on a trial-by-trial basis: target stimulus intensity (higher or lower), MOs activity, pre-stimulus eye movement, pupil diameter, running speed at target presentation, and pre-stimulus activity throughout the imaged cortex.

We computed a logistic regression model for each mouse (n=10, see Methods). Beta coefficients reflect predictor importance and the effect on behavior, with positive magnitude increasing the likelihood of the lick response (Fig. 5A, red points are significant within mouse). As expected, high-intensity trials were predictive of hit trials. Hit trials were also predicted by MOs activity, consistent with high choice probability and muscimol experiments (Figs. 3,4). Eye movement before target presentation increased the chance of detection failure (p=3.68×10^-3^, two-tailed t-test of beta coefficients). Interestingly, pupil diameter significantly predicted miss trials after correction for multiple comparisons (p=0.034, two-tailed t-test of beta coefficients). Run speed did not have a consistent effect across mice, although was significantly disadvantageous in one mouse (p=2.25×10^-8^, n=212 hit and 75 miss trials).

To assess model fit we computed d’ and prediction accuracy (see methods) for each mouse. All models had significantly higher d’ than the permuted-trial models (>95^th^ percentile, 1000 permutations, Fig. 5B). We reduced the model one predictor at a time (via permutation) to quantify how well each predictor explains mouse responses. Permutation of the MOs response variable had a large effect on d’ and prediction (Fig. 5B, “-MOs”, p=0.0034, paired t-test of d’ between permuted high intensity and permuted high intensity + MOs models). The model was still significantly better than chance when only eye movement was included (p=0.0102, paired t-test with permuted-trial model d’). Therefore, this analysis confirmed that MOs activity was indeed predictive of the lick response while revealing other variables with effects on hit rate.

### Widespread encoding of behavior and task variables

In the previous sections we found complex, overlapping representations of stimulus, response, arousal and running. We next sought to deconstruct activity across cortex into these component processes. Based upon published reports indicating that orofacial movements may have large explanatory power in predicting cortical activity in mice (Musall et al. 2019; Stringer et al. 2019), we selected a subset (n=5) of mice for which the videos included the nose, eye, mouth, and lower jaw (see Fig. 6B). We performed ridge regression within each session to explain cortical pixel activity on a trial-by-trial basis using ten predictors (Fig. 6A): 1) stimulus type (visual or auditory or none); 2) visual stimulus intensity (higher or lower); 3) visual trial response type (hit/miss/FA); 4) time of response; 5) time since last stimulus; 6) motion energy within the entire face video; 7) motion energy within an ROI centered on the whisker pad (see Fig. 6B); 8) speed of walking; 9) amplitude of eye movements; 10) pupil diameter (see Methods). Catch trials were random periods taken from the ITIs, so as to examine the rate of spontaneous licking and cortical activity. Trials with licking before 400 ms after the stimulus were omitted from the analysis.

Comparing the amplitude-time course of neural activity within regions of interest within MOs, S1, and V1 with the model fits revealed a remarkably good correspondence, indicating that the model was able to capture many of the variations in neural activity within different trial types (catch, visual, auditory; Fig. 6C). Examining the timing of bouts of licking, whisking, and walking revealed that these movements may be highly correlated with widespread neural activity (Fig. 6C). Maps of the total explained variance by the model revealed between approximately 35 to 55% explained variance throughout dorsal cortex, peaking in MOs and somatosensory cortical regions (Fig. 6D). By running the model with only one variable, we revealed the ability of that variable to explain the average variance of neural activity in the entire cortical image (Fig. 6E, red bars). Motion energy in the face video, or restricted to the whisker pad, were the two best predictors of the average neural activity changes (37+/-4.7%; 33+/-4.0%). This was followed by walk (25+/-4.4%), response choice (25+/-0.6%), eye movements (20+/-0.4%), response time (17+/-2%), light intensity (15+/-2.2%), stimulus type (10+/-1.8%), pupil diameter (8+/-1.8%), and time within the trial (5+/-0.8%) (mean+/-SD). By comparing the full model to reduced models generated by permuting one predictor at a time we calculated the unique explained variance of each predictor (Fig. 6E). Movement variables (walk, face movement, eye movement, whisking) again made significant contributions. Interestingly, global arousal, as indicated by pupil diameter, also made a significant unique contribution (Fig. 6E). Permuting all movement (video motion energy, whisk motion energy, walk, eye movement) along with pupil diameter revealed that these variables together contributed 23% (+/-1.7%) unique variance (not shown).

Next we examined the spatial distribution of β weights for each component of the model. Beta weight distributions for motion in the face video was concentrated in somatosensory and secondary motor (MOs) cortices (Fig. 6F), while whisk movement exhibited negative β weights concentrated in lower somatosensory cortex and motor cortical areas (Fig. 6G). Eye movement β weights were concentrated in visual cortical areas (Fig. 6H), while choice (Hit/Miss/FA) β weights were concentrated in MOs (Fig. 6I). These results reveal that it is possible to identify the spatial patterns of multiple behavioral and task-related representations across dorsal cortex, and that a large fraction of trial-to-trial variability and within trial activity can be explained by spontaneous or task related movements of the face (e.g. whiskers, licking, eye movements) and body (e.g. walking).

## Discussion

By monitoring widefield dorsal cortical activity, along with task relevant and/or spontaneous behavior, during the performance of a visual detection task, we were able to reveal patterns of cortical activity that are highly related to behavior of the animal (and vice versa). Our study revealed several general findings concerning this relationship. First, broad regions of the cortex are activated during performance of a simple Go/NoGo visual detection task, with earliest activation occurring in visual cortical areas, and more anterior regions of cortex becoming activated in relation to the lick response and associated orofacial movements. Second, activity in secondary motor cortex (MOs) was highly related to the generation of licks, either learned or spontaneous, and unilateral block of activity within this region resulted in a significant reduction in task performance (decreased appropriately timed licks and increased inappropriately timed licks). Third, predictive models of the neural activity revealed that the state of the animal (movement, arousal) explained a large fraction of within trial and trial-to-trial variation in neuronal activity. Finally, the behavioral choice of the animal (hit, miss, false alarm) explained a significant fraction of activity in the secondary motor cortical region (and vice versa). Taking performance, movement, and arousal variables into account, we were able to account for a significant fraction of dorsal cortical activity during the performance of the task.

### Secondary motor cortex and relation to performance of learned lick response tasks

In similarity with out present study, previous investigations have demonstrated a key role for an anterior cortical region, termed anteriolateral motor (ALM), premotor, or M2/MOs, in the performance of tasks involving learned lick responses (Allen et al. 2017; Komiyama et al. 2010; Guo et al. 2014; Li et al. 2016; Inagaki et al. 2019; Li et al. 2015; Makino et al. 2017; Goard et al. 2016). Widefield imaging during performance of lick-response tasks has revealed strong activation of M2/ALM (Allen et al. 2017; Makino et al. 2017), and bilateral block of this cortical region results in a large decrease in the ability of the mouse to perform the task accurately (Allen et al. 2017; Komiyama et al. 2010; Li et al. 2016; Goard et al. 2016). We found using unilateral block of M2/ALM that hit trials are associated with broad activation of the non-frontal cortical regions (Fig. 4D), presumably owing to the ability of the contralateral M2/ALM area, which remains active (Fig. 4D), to contribute to performance of the task (Li et al. 2016). This broad cortical activation was not seen on miss trials (although activation of visual cortex by the stimulus remained) (Fig. 3), suggesting that previous observations of a lack of activity in cortical regions following bilateral inhibition of M2/ALM was the result of subsequent non-performance of the task (Allen et al. 2017).

Recording neural activity, either optically or electrophysiologically, in M2/ALM during performance of learned lick responses reveals neuronal activity that precedes and predicts the performance of licks (Komiyama et al. 2010; Li et al. 2016; Inagaki et al. 2019, 2018; Chen et al. 2017; Guo et al. 2014; Erlich, Bialek, and Brody 2011; Zagha, Ge, and McCormick 2015). While recording and inactivation studies indicate that M2/ALM is critical to the performance of the learned motor response in mice, it is unclear at present whether or not this critical role is limited to learned movements. Stimulation of M2 in naive rodents can initiate licks (Travers, Dinardo, and Karimnamazi 1997), and we found M2 (MOs) to be strongly activated even during spontaneous licks (Fig. 4A,B), suggesting that M2 is important for the cortical control of both learned and unlearned lick movements.

### Relationship between movement, state, cortical activity and task performance

The behavioral state of waking mice, including movement and arousal, is in constant flux (reviewed in (McGinley et al. 2015; Poulet and Crochet 2018)). These fluctuations in state, which can occur on a sub-second time course, can significantly impact performance on sensory detection or discrimination tasks (McGinley et al. 2015; McGinley, David, and McCormick 2015; Poulet and Crochet 2018). Changes in level of arousal, as indicated by pupil diameter, is an important factor in controlling brain state, from the activity patterns of single neurons (e.g. burst versus tonic), to local cellular interactions, to the broad operation of cortical and subcortical circuits (McCormick, McGinley, and Salkoff 2015). Likewise, movement, which is highly interwoven with arousal, can also have a powerful impact on the activity of cortical networks, not only in somatosensory and motor cortical areas (Poulet and Crochet 2018; Petersen 2007), but also in other primary sensory pathways such as visual and auditory cortices (Niell and Stryker 2010; McGinley et al. 2015; McGinley, David, and McCormick 2015; Stringer et al. 2019; Musall et al. 2019; Shimaoka, Harris, and Carandini 2018). Indeed, recent reports indicate that subcomponents of facial motion, either spontaneous or during the performance of a task, is able to explain a significant fraction of dorsal cortical activity, as monitored either through optical, or electrical, physiology (Stringer et al. 2019; Musall et al. 2019). In agreement with these studies, our current findings reveal that linear models of widefield cortical activity are able to explain a significant component (up to 55% of the variance) of this activity, not only in somatosensory and motor cortical regions, but also in primary visual cortex and other adjacent cortical areas (e.g. retrosplenial cortex; Fig. 6D). Movement forms the largest component of explained variance in our study, as in previous investigations (Stringer et al. 2019; Musall et al. 2019), indicating that a significant fraction of trial-to-trial variation in neural responses is the result of changes in behavioral state between trials. This high level of explained variance is the result, in part, of examining widefield activity, which blurs together the spatial-temporal patterns of neuronal cell bodies and processes within the imaged region. Examining facial/body movements and task variables is also able to explain a significant amount of neuronal action potential activity within the cortex and subcortical regions (Musall et al. 2019; Stringer et al. 2019), although this explained variance is lower than for the general features of widefield activity (Musall et al. 2019).

Examination of the interaction of spontaneous, or ongoing, cortical activity and sensory evoked responses indicate that alterations in the state of activity can have strong effects on the amplitude, timing, and reliability of the sensory evoked response (McGinley et al. 2015; McGinley, David, and McCormick 2015; Poulet and Crochet 2018). Similarly, we observed here that changes in pre-trial activity are associated with trial performance, with increased activity in somatosensory and visual cortical areas being more prevalent on miss trials (Fig. S9).

### Possible Mechanisms of State and Performance Dependent Variations in Cortical Activity

The complex patterns of widefield cortical activity occurring in behaving animals performing a simple Go/NoGo visual detection task result from ongoing changes in a wide variety of variables, from shifts in arousal and task engagement, to task related and task-unrelated movements (e.g. licking, eye movements, walking), to the operation of neural circuits critical for detection of the stimulus and performance of the motor response. Many of these variables are inter-related. Movement, for example, is strongly influenced by the sleep-wake cycle, arousal, and attention (and vice versa).

Ascending cholinergic and noradrenergic pathways are both tonically activated by slow increases in arousal (McCormick 1992; McCormick, McGinley, and Salkoff 2015), as well as phasically activated by sudden changes in arousal, movement, or task engagement (Reimer et al. 2016; Eggermann et al. 2014; Pinto et al. 2013; A. M. Lee et al. 2014; Nelson and Mooney 2016). These modulatory transmitter pathways can have both rapid (10s of msec) and slow (seconds) effects on activity in cortical and thalamocortical networks (Muñoz and Rudy 2014; Zagha and McCormick 2014). For example, the activation of nicotinic receptors by acetylcholine may rapidly activate layer 1 VIP-containing interneurons, resulting in the inhibition of somatostatin containing inhibitory cells and the disinhibition of pyramidal neurons (Pi et al. 2013; Zhang et al. 2014; S. Lee et al. 2013; Hangya et al. 2014), thereby participating in the rapid changes associated with microarousals, or short duration movements (McGinley et al. 2015; Drew, Winder, and Zhang 2018). Through the activation of metabotropic receptors, noradrenergic and cholinergic systems can also have broad effects on the patterns of cortical and thalamocortical networks by altering neuronal excitability (e.g. depolarization) and neuronal firing pattern, among other effects (Lee and Dan 2012; McCormick 1992; Zagha and McCormick 2014). While the temporal possibilities for neuromodulation have been detailed, the spatial specificity of known is these modulatory influences are not yet known. While cholinergic systems may have the anatomical specificity to influence small regions of the cerebral cortex, the widespread connections of individual noradrenergic axons suggest a more spatially broad influence (Muñoz and Rudy 2014; Kebschull et al. 2016; Schwarz et al. 2015). The activity of many cortical cholinergic and noradrenergic axons is strongly correlated with pupil diameter, a general proxy for arousal (Reimer et al. 2016), suggesting a broad modulation of cortical areas. While measurements of arousal (as indicated by pupil diameter) can facilitate the explanation of trial-to-trial variance in overall neuronal gain and task performance (McGinley et al. 2015; McGinley, David, and McCormick 2015), general arousal cannot account for the rapid alterations in complex patterns of cortical activity we observed in awake, behaving mice, indicating the involvement of more spatially and temporally precise mechanisms.

The dorsal cortex of the mouse is dominated by motor, somatosensory, and visual cortical areas. Movement, and its associated somatosensory feedback, is expected to result in strong activation of these cortical regions. In addition to the somatosensory and motor cortical areas, the visual cortex of the mouse is also strongly activated by movement (Niell and Stryker 2010; Khan and Hofer 2018). The representation of the face, and its components (e.g. whiskers, tongue, lips, nose, eyes, etc.), occupies a significant fraction of somatosensory and motor cortical areas. In addition, movements of the face are often coordinated with movements of the forelimbs and body (e.g. feeding, grooming). By monitoring even the gross movements of the face (e.g. whisker pad motion energy) and body (e.g. walking), a significant fraction of the activity of the dorsal cortex, as observed at the widefield or even action potential level, can be predicted (Fig. 6; (Stringer et al. 2019; Musall et al. 2019)). These results indicate that dorsal cortical neural signals, integrated over local space (on the order of approximately 100 um) and time (100s of msec), are strongly related to either the performance of, or sensory feedback from, facial and bodily movements. The highly interconnected nature of the cerebral cortex will further allow for signals in one area (e.g. M2/MOs) to influence the activity in others, including primary sensory cortical regions (Nelson et al. 2013; Schneider, Nelson, and Mooney 2014; Schneider and Mooney 2018; Leinweber et al. 2017; Zagha et al. 2013; Makino et al. 2017; Khan and Hofer 2018; Felleman and Van Essen 1991). Presumably, the broad impact of movement on a broad range of cortical areas, including sensory regions, results from the need to take into account self motion (Schneider and Mooney 2018; Leinweber et al. 2017) in a manner consistent with predictive coding (Keller and Mrsic-Flogel 2018), since movement and sensory coding are strongly coupled together during natural behavior. Thus, given the importance, and complexities, of orofacial movements in the normal behavior of mice, and specifically in the performance of learned tasks in which the response is a lick, it is perhaps expected that broad regions of the dorsal cortex would be activated in association with performance of the task and that movements of the face and body may be the most salient features to explain this activity (Musall et al. 2019; Stringer et al. 2019).

Which parts of the cerebral cortex are critical for performance of learned lick responses to sensory stimuli? Previous investigations, which have surveyed a broad range of cortical areas through inactivation, have revealed inactivation of two cortical regions to be effective in blocking task performance: 1) the primary sensory cortex (e.g. somatosensory) involved in detection; 2) premotor (ALM/M2) cortex (Allen et al. 2017; Guo et al. 2014; Li et al. 2016). Our current observation of significantly decreased performance of learned licks to a visual stimulus following unilateral inactivation of M2/ALM (Fig. 4E) confirms these earlier results. The increased rate of inappropriately timed licks (i.e. false alarms; Fig. 4E) indicates that this decrease in performance is not simply a reflection of the inability to perform licks, but rather suggests difficulties with generating maximally efficient responses. Monitoring the activity of neurons in M2/ALM, either optically or electrophysiologically, reveals preparatory activity that anticipates, and potentially encodes, the performance of the upcoming response (Guo et al. 2014; Chen et al. 2017; Komiyama et al. 2010; Li et al. 2016, 2015; Inagaki et al. 2018; Allen et al. 2017; Erlich, Bialek, and Brody 2011). This preparatory activity has the hallmark of motor planning that utilizes discrete attractor dynamics within an interconnected network of cells (Inagaki et al. 2019). Whether or not this network of neurons is actively involved in evidence accumulation for detection of the stimulus, or more closely related to preparation for the upcoming movement execution, remains to be determined. Even so, it is becoming increasingly clear that only through careful monitoring of variations in both spontaneous and task-related behavior and state (e.g. arousal, task engagement) will the neural signals underlying signal detection, decision, and response implementation be revealed.

## Acknowledgements

We would like to thank Gregg Castellucci, Jake Lister, and members of Dr. Michael Crair’s lab for helpful feedback throughout the project; Doug Storace for thoughtful conversations regarding controlling for hemodynamic and motion artifacts; Alex Kwan for feedback on the manuscript; and Trevor Stavropoulos and Tony Desimone for technical assistance.

## Supplement and Supplementary Figures

### Spontaneous and hit-related running/arousal changes

Running before the target was sometimes followed by significant slowing, or stopping, by 0.5-0.7 seconds post-stimulus (Fig. S1). Mice which had been sitting prior to the target often initiated running on hit trials. In the session shown in Figure S1A, initiation of wheel movement (>0.5 cm/s) occurred 423 ms (±108, n=10 mice) after presentation of the lower-intensity target. Movement occurred significantly faster after the higher-intensity target (317 ±95 ms, p=1.8×10^-4^, paired t-test, n=10 mice). Similarly, lick response time was significantly faster after the higher-intensity compared to lower-intensity target (2.5×10^-4^, paired t-test, n=10 mice). Treadmill speed was often highly stereotyped within sessions. However, we observed different strategies for different mice, and even for the same mouse in different sessions (Fig. S1A,B).

We wondered whether changes in pupil diameter reflected changes in locomotion. Cross-correlation of run speed and pupil diameter shows high correlation between the two (Fig. S1C, maximum correlation 0.473 ±0.103, n=10 mice). Correlation was highest at −0.510 sec (±0.208) lag, suggesting pupil diameter lags, on average, running. We confirmed this by looking at simultaneously recorded treadmill speed and pupil diameter locked to either target presentation on hit trials (Fig. S1A) or spontaneous transitions from still to running (Fig. S1B). In both cases running changes preceded pupil diameter changes. The lag between changes in running speed and pupil diameter may largely reflect the fact that pupil diameter changes lag sympathetic/parasympathetic activity by approximately 0.5-0.75 seconds.

**Figure S1.**
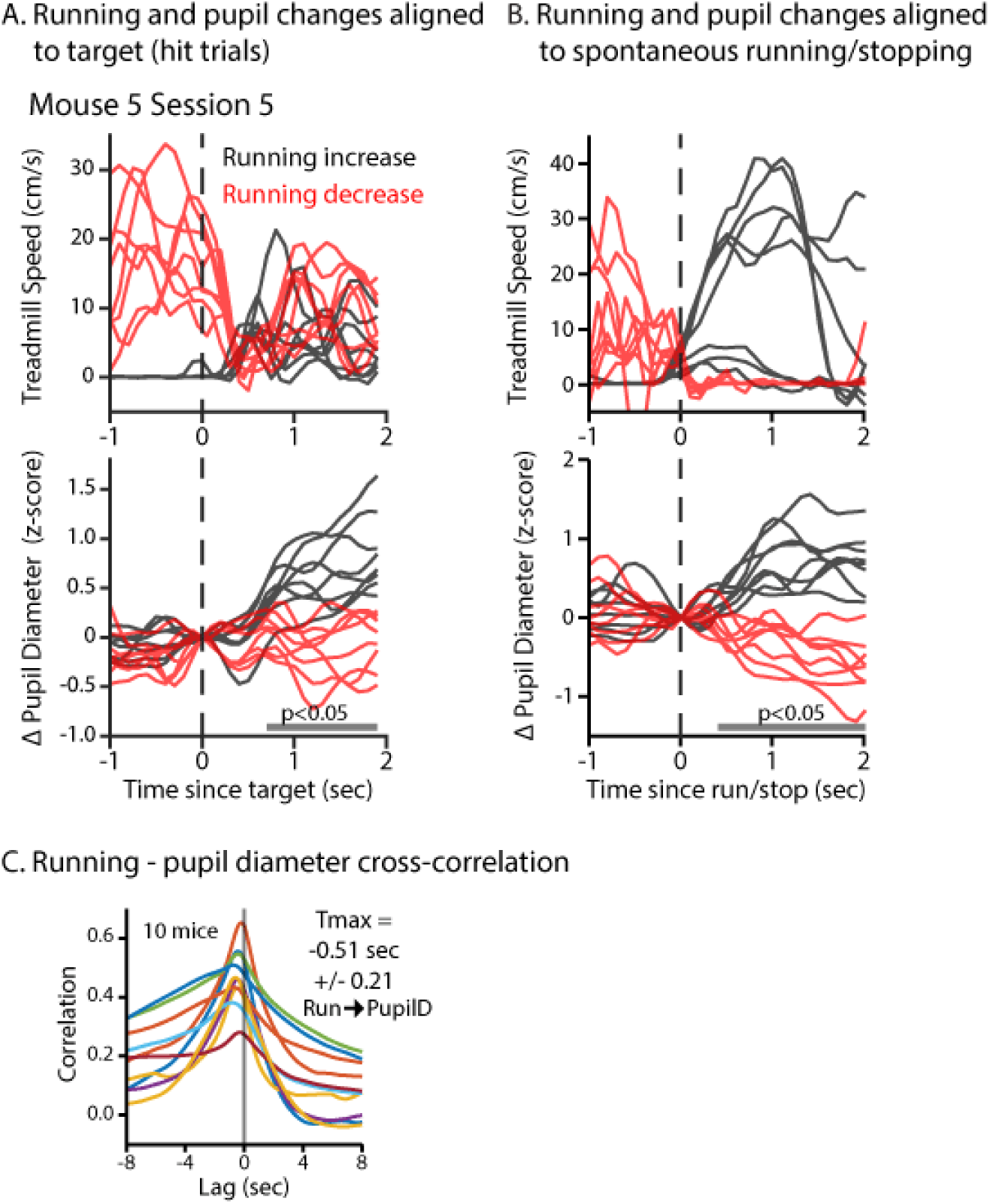
Spontaneous and target-related running/arousal changes. **A**, Running (top) and corresponding pupil (bottom) traces on example hit trials. Trials have been color coded as increased-running (black) or decreased-running (red). Note all increased-running trials result in higher pupil diameter than the decreased-running trials. **B**, Running (top) and corresponding pupil (bottom) traces aligned to either the time of spontaneous running initiation (black) or spontaneous running cessation (red). **C**, Cross-correlogram showing high correlation between running velocity and pupil diameter (n=10 mice). Pupil increases lag behind running increases on average.

**Figure S2.**
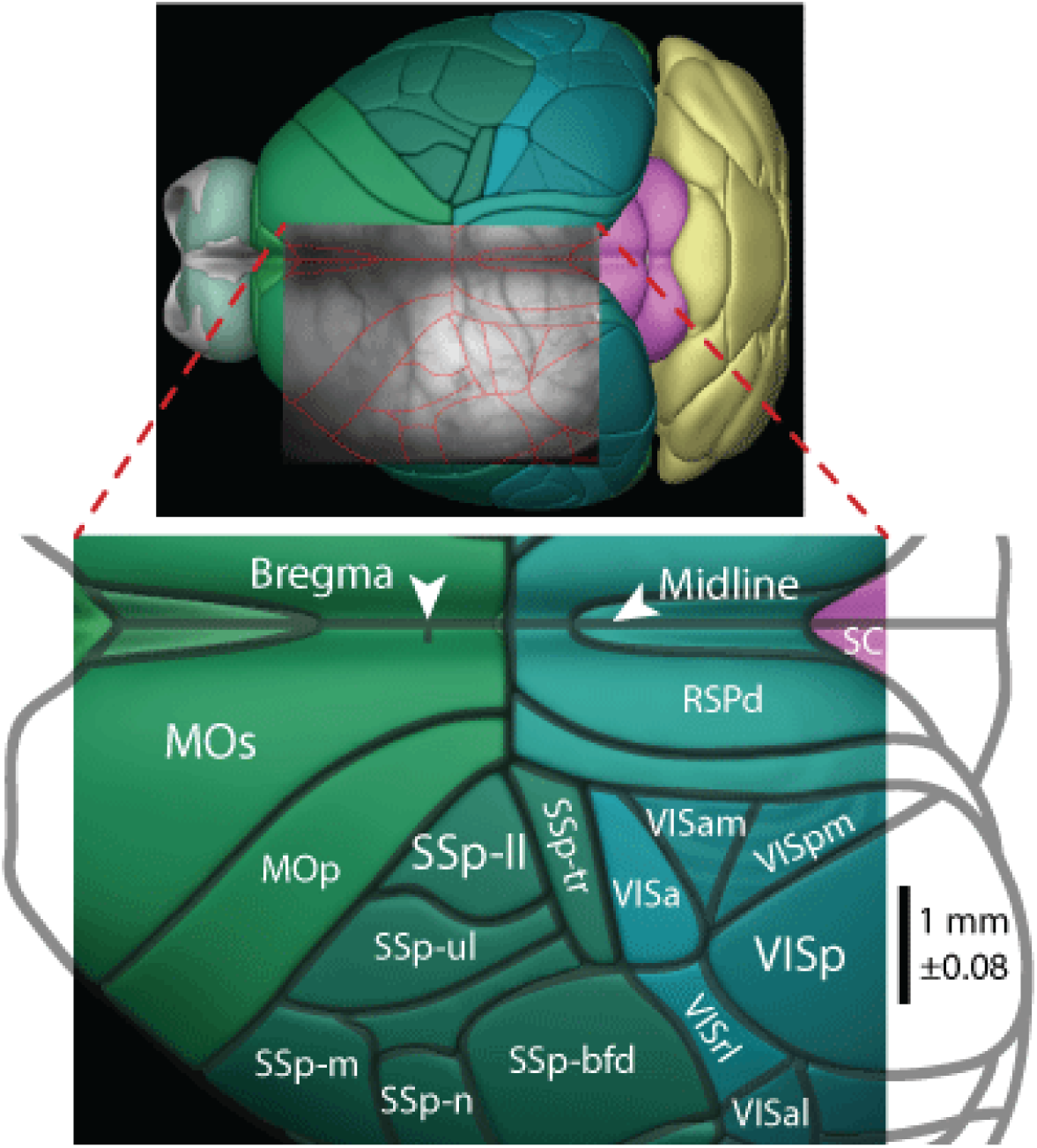
Wide view of dorsal cortical surface that was imaged during task performance. Top: 3D brain model from the Allen Mouse Common Coordinate Framework showing dorsal view. We imaged the cortical hemisphere contralateral to a visual stimulus associated with water reward. An image from each mouse was manually aligned using bregma and lambda, then subsequently confirmed using functional activity (example image from one session shown overlaid). Bottom: dorsal cortex with lines delineating regions from the corresponding raw image above. Images of cortex from each mouse were (rigidly) stretched and rotated to match the atlas. Scale bar indicates the average length of 1.0 (±0.08) mm. Abbreviations are from the atlas mentioned above, but defined here (regions of interest emboldened): MOp, primary motor; **MOs, secondary motor**; SC, superior colliculus (sub-cortical); RSPd, dorsal part retrosplenial; SSp-bfd, primary somatosensory barrel-field; **SSp-ll, primary somatosensory lower-leg**; SSp-m, primary somatosensory mouth; SSp-n, primary somatosensory nose; SSp-tr, primary somatosensory trunk; SSp-ul, primary somatosensory upper-leg; VISa, anterior visual (also known as PTLp, posterior parietal); VISal, anterolateral visual; VISam, anteromedial visual; VISl, lateral visual; **VISp, primary visual**; VISpm, posteromedial visual; VISrl, rostrolateral visual (also known as PTLp, posterior parietal).

**Figure S3.**
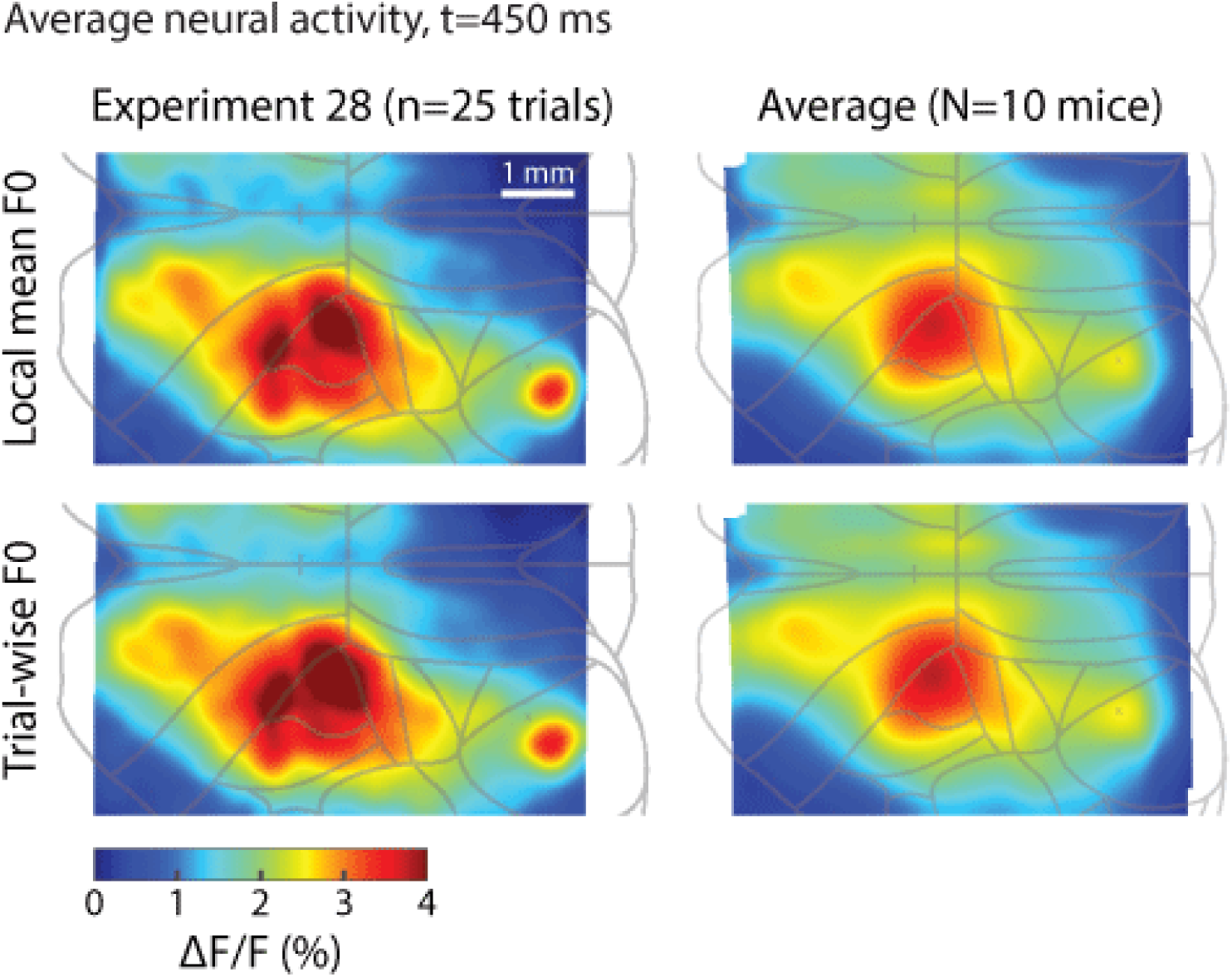
Comparison of normalization method where ΔF/F = (F_i_-F0)/F0. F0 in this paper was calculated by averaging the fluorescence per pixel within ± 4 minutes (local mean method). This allowed us to compute a continuous readout of neural activity. In contrast, standard ΔF/F calculates F0 from the pre-stimulus activity of each trial (trial-wise method). This figure shows the similarity of the two methods.

**Figure S4.**
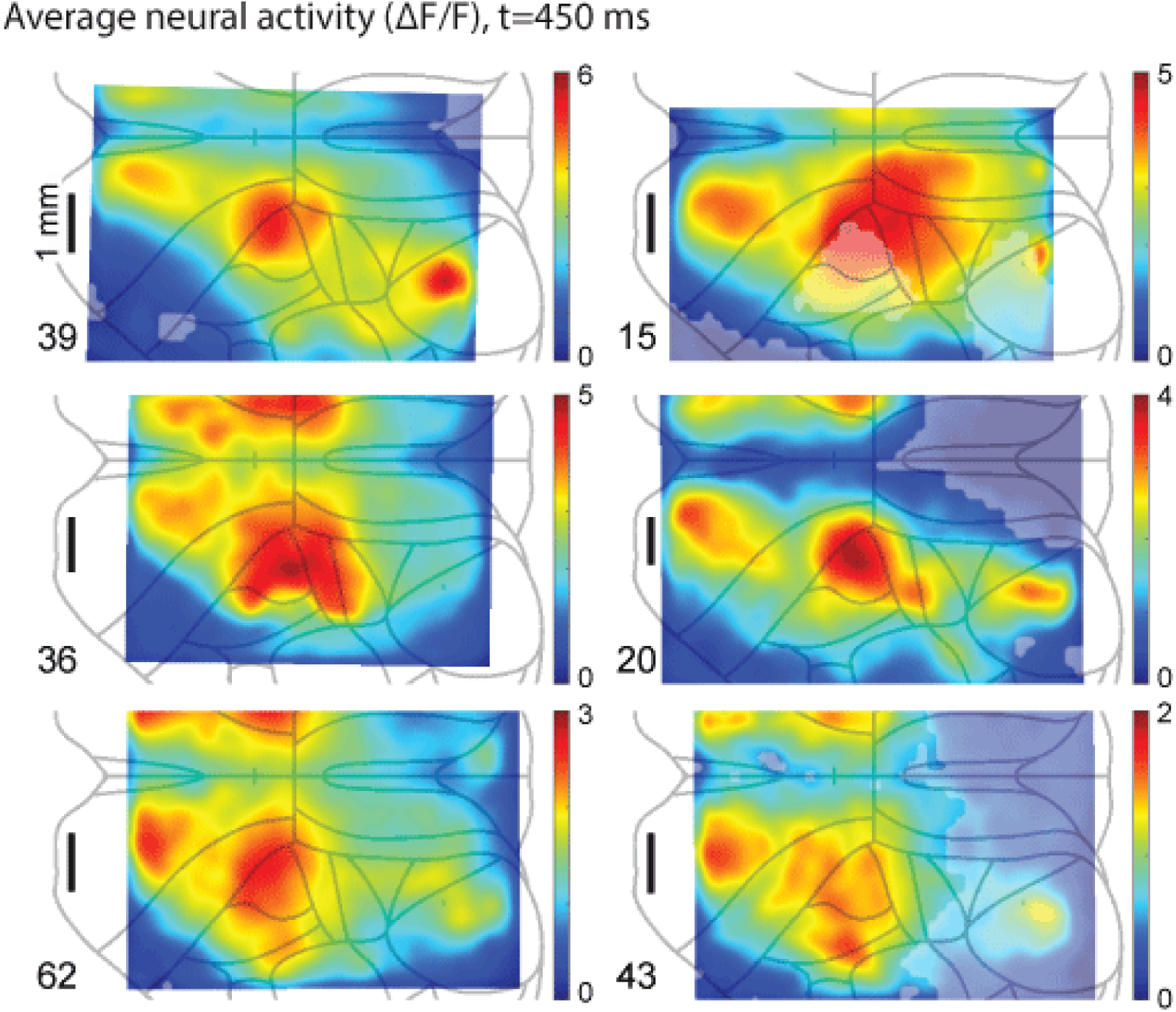
Additional examples of post-stimulus activity (n=6 mice). Each map is the average neural activity 450 ms after the higher-intensity target computed from one session of different mice (similar to Fig. 2B). The number of trials averaged is indicated in the bottom left of each map. Trials with response time faster than 500 ms were omitted from the analysis. Opaque pixels indicate significantly large clusters of pixels with p<0.05 (p<0.05, permutation test between pre and post-stimulus activity). Note similar activation of frontal and parietal cortices yet variation in precise spatial structure of maps. Sources of animal-to-animal variability in fluorescence intensity include effectiveness of the skull-clearing procedure, individual differences in expression levels and individual differences in neural/behavioral strategies for task completion. Since our analyses focus primarily on the patterns of activation rather than their magnitude, we do not believe this to significantly obscure our main findings. See supplemental movie for real time examples of activity patterns.

### GCaMP6s fluorescence reflects regional neural activity

We investigated the spatio-temporal relationship of the GCaMP6s signal to ensure that widespread activity after target detection was not a result of movement artifact or spatial scatter through the skull. The relationship between neural activity and GCaMP6s fluorescence has been described *in vivo* on a single-cell basis (Chen et al. 2013), however the spatiotemporal properties of this calcium indicator are still being investigated at the scale of neural populations through a trans-cranial window. To this end we performed simultaneous electrophysiology and wide-field imaging experiments. An extracellular electrode was placed in the parietal cortex of an anesthetized mouse (ketamine/xylazine). The anesthetized state is characterized by the slow oscillation (0.1-4 Hz), a periodic increase in neural firing followed by quiescence ((Otchy et al. 2015; Sirotin and Das 2009; Steriade, Nunez, and Amzica 1993). Band-pass filtering of the local field potential (LFP) revealed periods where both gamma activity and neuronal firing rates were high (Fig. S5A). An overlay of the fluorescence (ΔF/F) from the cortical region surrounding the electrode tip shows a high correlation between neuronal firing and GCaMP6s activity. Fluorescence started to increase almost immediately after neuronal firing started. This signal increased until after firing stopped and then decayed until the next phase of neuronal firing. Cross-correlation analysis confirmed that fluorescence was highly correlated with gamma activity (r=0.726). Maximal ΔF/F lagged behind peak gamma amplitude by 0.4 seconds, although this lag may be more indicative of the duration of the active phase of the slow oscillation than fluorophore dynamics. These analyses indicate that the Ca^2+^ signal in our preparation highly correlates with spiking activity, possibly reflecting the integration of neuronal firing.

We next sought to understand the spatial extent of fluorescence signals. Scattering of fluorescent light, either through superficial cortex or bone, may cause active cortical regions to appear larger than they actually are. Neural activity in trained mice appeared widespread (Fig. 2), and therefore we wondered if this was truly a result of widespread cortical activity or whether it was an artifact of spatial scattering. We therefore habituated two naïve mice to head-fixation on the treadmill and recorded neural responses to spontaneous 50 ms flashes similar to the target stimulus in the visual detection paradigm. Visual responses were apparent on single trials (Fig. S5B). Inspection of the activation probability map revealed a more focal and reduced response compared with the pre-lick neural activity of trained mice (Fig. S5C, compare with Fig. 2C,F). On average, activity arose in VISp first followed shortly after by retrosplenial cortex and spread to surrounding areas (Fig. S5D). These experiments in naïve mice provided an understanding of the spatial extent and reliability of unconditioned visual responses when imaging trans-cranially. Moreover, they demonstrate that light scattering cannot account for the widespread optical signals we observe during behavior.

We sought to further understand the extent of neural correlation. By “seeding” an arbitrary pixel and calculating the correlation of ΔF/F with every other pixel we can observe how rapidly correlation decays across the surface of cortex (Fig. S5E). Correlation was widespread in the anesthetized state, regardless of seed location. This was not surprising given the high degree of spatial synchrony during the slow oscillation (Massimini et al. 2004). In contrast, correlation structure during the awake, task-engaged state (Fig. 1) was more spatially refined. We observed that correlation generally decreased monotonically as distance from the seeded pixel increases. This was not the case for some seeds such as secondary motor cortex which was highly correlated with a somatosensory region representing the nose and mouth. We observed local gradients as high as 1.2 R^2^/mm. Assuming linearity, this implies two uncorrelated brain regions ≥0.84 mm apart have independent fluorescence signals.

**Figure S5.**
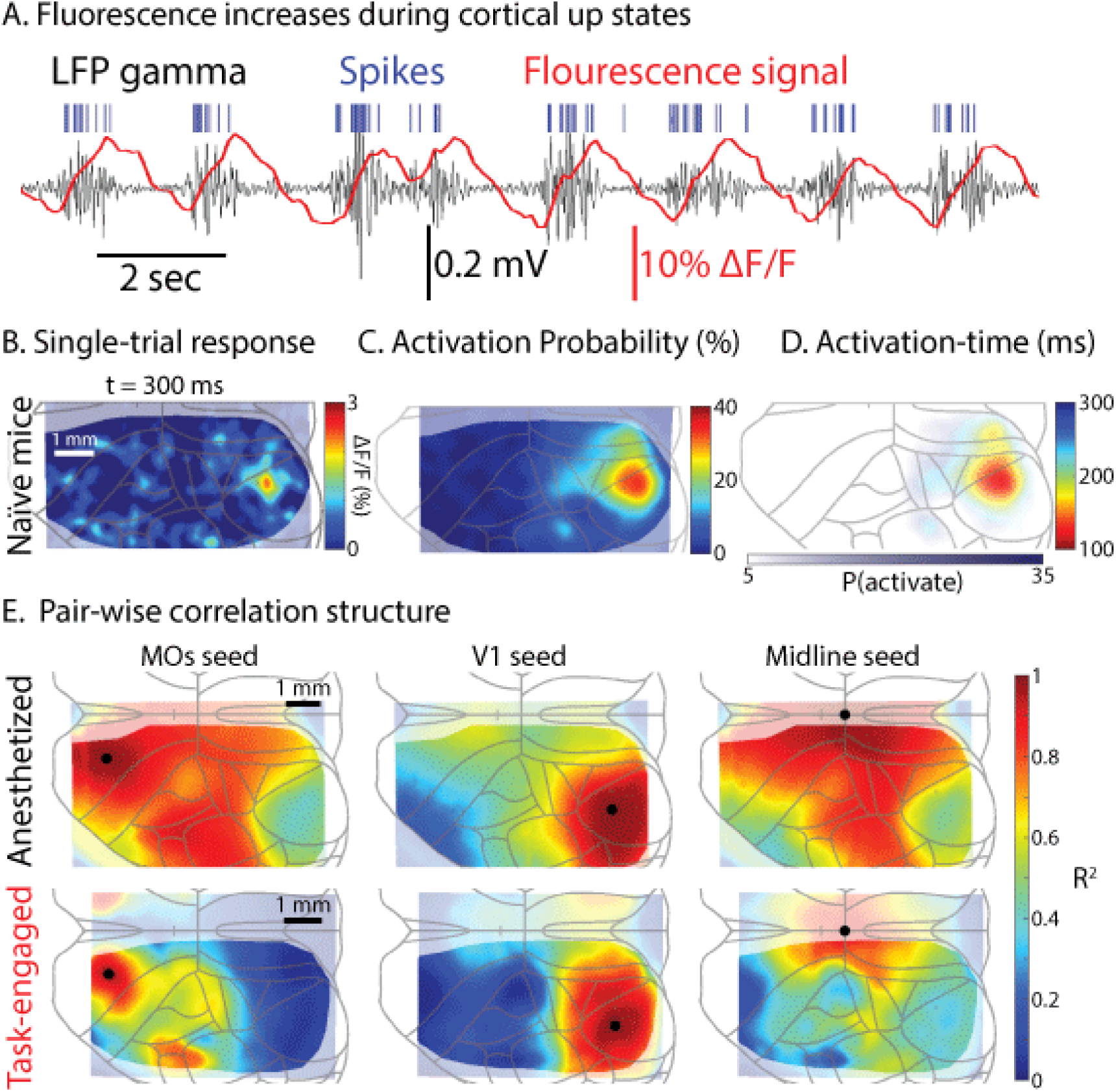
GCaMP6s fluorescence reflects regional neural activity. **A**, Simultaneously measured local-field potential (LFP band-pass filtered 20-80 Hz) and fluorescence (ΔF/F) measured in the region of the electrode (parietal cortex). Recordings are from a ketamine-xylazine anesthetized mouse. Note the high correlation between gamma, spiking, and fluorescence. **B**, Single-trial visual response from a naïve mouse. **C**, Activation probability after t = 450 ms, average of two naïve mice and six sessions. Note the restricted spatial extent in comparison to Fig. 2C,F. **D**, Average activation-time map, same mice as C. **E**, Pair-wise correlation structure in anesthetized and task-engaged states. Each column has a different “seed” pixel (black dot) which was cross-correlated with all other pixels. Correlation was generally more spatially restricted in the task-engaged state, with maximum gradients of 1.2 R^2^/mm observed, suggesting uncorrelated cortical regions as close as 0.84 mm apart may appear to have independent fluorescence signals.

### Hemodynamic and motion artifacts do not account for widespread fluorescence changes after target perception

Increased fluorescence after target presentation could be caused by motion artifacts when the mouse shifts in preparation to lick. Fluorescence may also change due to hemodynamic effects: either sniffing-induced changes in blood pressure or when blood vessels dilate/contract in response to neuronal activity (Wekselblatt et al. 2016) or in anticipation of salient information (Otchy et al. 2015; Sirotin and Das 2009). We measured changes in fluorescence unrelated to GCaMP6s in two ways. First we measured reflected light in a GCaMP6s negative mouse which had been trained in the task. Reflectance changes in the GCaMP6s negative mouse after target presentation were much smaller than fluorescence changes in all GCaMP6s positive mice (Fig. S6A left panels, three areas of interest). A map of average increase in ΔF/F revealed the largest changes to be around the midline (Fig. S6A, right panel). Experiments in an additional two mice yielded similar results.

Next we measured motion and hemodynamic signals by recording under green (530 nm) light. GCaMP6s does not fluoresce at this wavelength, however absorption and reflectance should be similar to blue light by other sources (Wekselblatt et al. 2016). Changes in reflectance were normalized in the same way to produce ΔF/F. Green reflectance increases were small and comparable to those measured under blue light for the GCaMP6s negative mouse (Fig. S6B, middle panel). After recording several trials under green light we switched to blue light for the remainder of the session. This allowed us to subtract green light changes from those recorded under blue light (Fig. S64B, left panel). We computed the difference between the two light conditions using a permutation test (Nichols and Holmes 2002)(see methods). The difference in ΔF/F between these light conditions contained a cluster of pixels (each pixel with p<0.05) significantly larger than expected by the null hypothesis (p<0.001, Fig. S6B right panel, opaque pixel cluster). Two other mice recorded under both blue and green light had similar results (data not shown). Thus, even after correction for light changes unrelated to GCaMP6s fluorescence a widespread fluorescence increase was present. Experiments with GCaMP6s negative mice and GCaMP6s positive mice under green light elucidate the magnitude and spatial patterning of non-GCaMP6s related signals. Both these studies suggest that, while non-GCaMP6s signals are not negligible, they do not account for the spatial extent of activity after target detection and contribute only a minority of the signal obtained with GCaMP6s imaging (Fig. 2).

**Figure S6.**
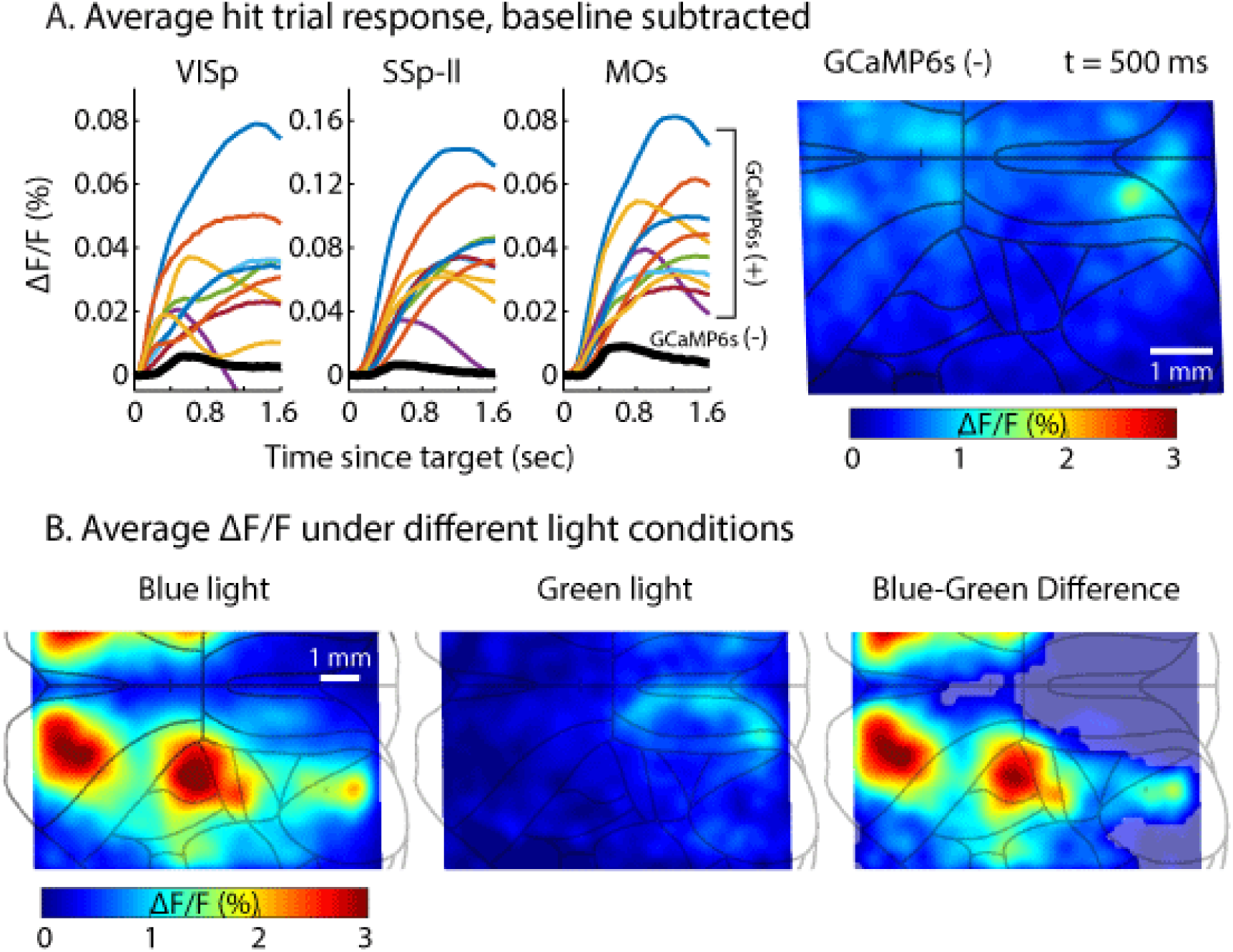
Hemodynamic and motion artifacts do not account for widespread fluorescence changes. **A**, Left: average (high-intensity) hit trial responses from all GCaMP6s (+) mice (n=10) and one GCaMP6s (-) mouse (black trace). Right: Average hit trial response from a GCaMP6s (-) mouse across cortex. The largest reflectance changes were around the midline, but were considerably smaller than fluorescence changes in GCaMP6s (+) mice. **B**, Average ΔF/F under different light conditions. A GCaMP6s (+) mouse was recorded under blue light (left) and green light (middle). The green light does not excite the GCaMP6s fluorophore but indicates changes in blood flow or sample movement. Several trials were recorded under each light condition. A permutation test revealed the difference (right) had a significantly large cluster (p<0.001), suggesting non-GCaMP6s related changes in ΔF/F do not account for widespread activity after target perception. Both control experiments (blue light stimulation in GCaMP6(-) mice and green light stimulation in GCaMP6(+) mice) were replicated with similar results for a total of n=3 mice each.

**Figure S7.**
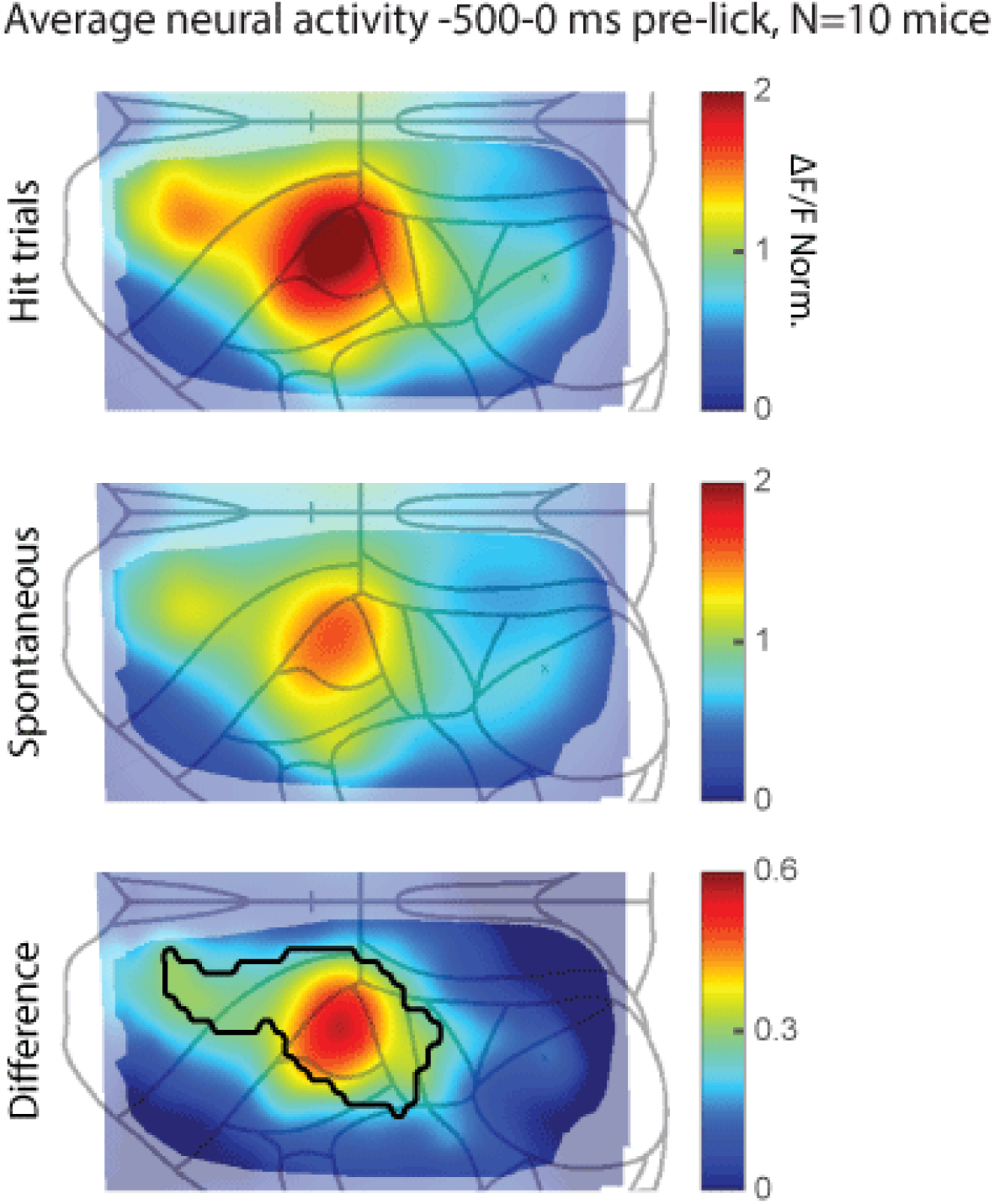
Neural activity leading up to licking. Average activity from 0 to 500 ms pre-lick in hit trials (top) and spontaneous licks during inter-trial intervals (middle) averaged across 10 mice. Bottom panel shows the difference between the two maps and a region that was significantly different (outlined in black, each pixel p<0.05) with size larger than expected from the null distribution (p=0.02).

### Somatosensory lower-leg region is activated in association with both running and licking

We sought to further characterize a cortical region exhibiting both sensory and choice-related activity (Figs. 2,3): the lower-leg region of primary somatosensory cortex (SSp-ll). Inspection of SSp-ll activity alongside treadmill speed revealed a high correlation between the two (Fig. S8A), presumably resulting from somatosensory and motor aspects of running. We aligned SSp-ll activity to spontaneous running initiations, and observed prominent activity increases (Fig. S8B). Similarly, transitions from running to still resulted in prominent decreases in SSp-ll activity (Fig. S8C). We cross-correlated cortical activity in every region with a vector indicating the still or running state (Fig. S8D). Running state accounted for ∼40% of variance in SSp-ll. Pupil diameter (a proxy of arousal, which is correlated with running - Fig. S1) accounted for ∼20%of variance in SSp-ll (Fig. S8E).

Closer inspection of SSp-ll revealed moments where activity did not match running speed (Fig. S8A, green arrows), but increased during licking regardless of running speed. To systematically test whether the relationship between SSp-ll and running changes during the lick response, we compared activity during hit trials with that during the inter-trial-interval (ITI) when similar running occurred (see methods). We first inspected activity during increased running (examples of running, S8Fi; associated activity, S8Fii). Target trials when the mouse was stationary at stimulus onset resulted in running increases with reliable onset times (Fig. S8Fi, red traces; also see Fig. S1). After finding spontaneous running transitions, we aligned these to the onset of running during hit trials. The activity associated with increased running was remarkably similar between hit trials and spontaneous events during the ITI (S8Fii, green arrow denotes 400 ms post-stimulus).

This was not the case for decreased running. We compared hit trial and ITI activity in SSp-ll, this time for decreased running (Fig. S8Fiii). While spontaneous run cessation during the ITI was often associated with decreases in activity (Fig. S8C, Fiv grey traces), hit trials with decreased running exhibited rapidly rising neuronal SSp-ll activity time-locked to the stimulus (Fig. S8Fiv red traces). We computed SSp-ll activity associated with increased/decreased running during hit-trials/ITIs for each mouse (Fig. S8Fv). Differences in running had a large effect on activity during the ITI (p=1.65×10-5, n=10, paired t-test). However differences in running during hit trials did not significantly affect activity. Together, these analyses suggest SSp-ll is activated by both running and performance of licks (either spontaneous or on hit trials).

**Figure S8:**
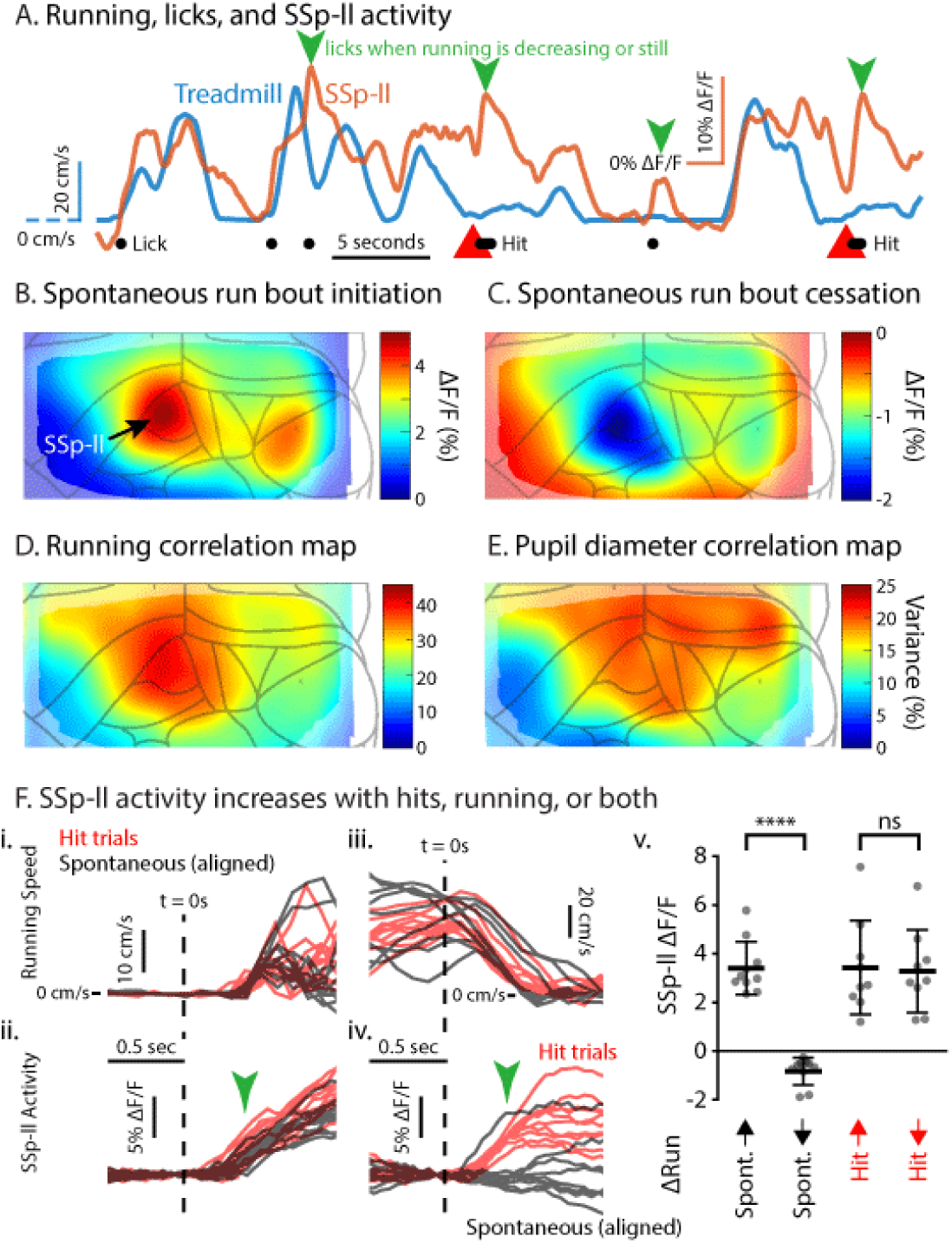
Somatosensory lower-leg region (SSp-ll) exhibits activity that is related to both running and licking, either spontaneous or during trials. **A**, Example of SSp-ll activity (orange) and mouse running speed (blue) alongside task variables (licking, black dots; red triangles: target). Note correlation between SSp-ll activity and run speed, however activity also increases during licking even when running speed is decreasing or not increasing strongly (green arrows). **B**, Average neural activity immediately after transitions from still to walking showing high activity in SSp-ll (n=10 mice). **C**, Same as B, but for transitions from walking to still. **D**, Neural variance explained by running. **E**, Neural variance explained by pupil diameter (which is correlated with running; see Fig. S1). **F**, SSp-ll activity increases with hits, running, or both. **i**, Running increases on hit trials (red) or during spontaneous transitions (black) for an example session. Spontaneous running traces have been aligned in time to match the onset of running during hit trials. **ii**, SSp-ll activity corresponding to the running traces in i showing similarity of activity at 400 ms (green arrow). **iii**, Running decreases on hit trials (red) or during spontaneous transitions (black). Traces have been aligned as in i. **iv**, SSp-ll activity corresponding to the running traces in iii showing difference of activity at 400 ms (green arrow). **v**, Summary of SSp-ll activity for 10 mice. Activity during spontaneous running and slowing has opposite magnitude, however SSp-ll activity remains high during lick responses (hit trials) even with decreases in running.

**Figure S9.**
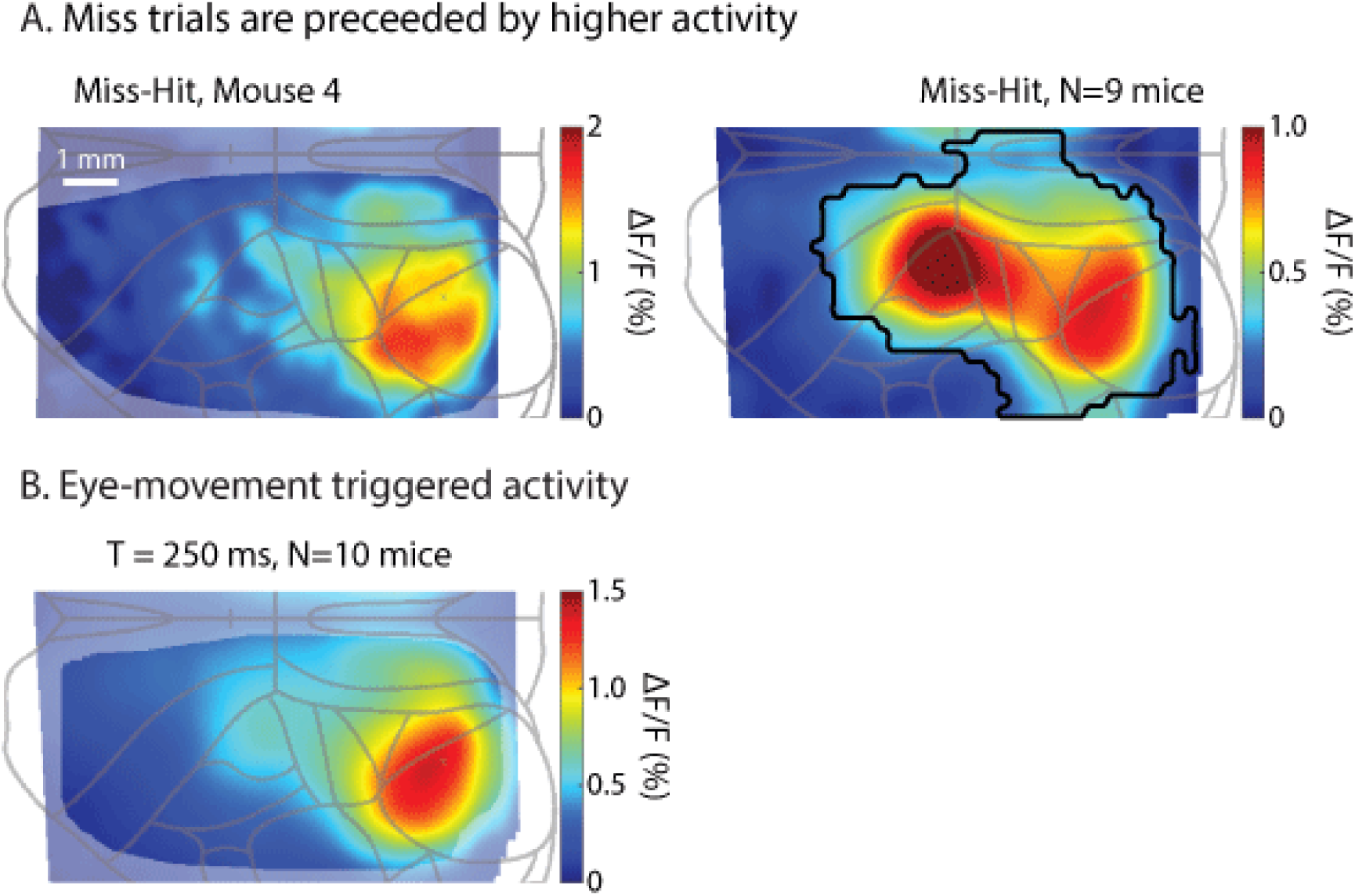
Pre-stimulus activity and eye-movement are possible sources of performance variability. **A**, Pre-stimulus activity was higher on miss trials than hit trials on average. Left: single session which had significant pixels (p<0.05) in occipital cortex but was not significant for cluster size (p=0.072). Right: Comparison of the average hit/miss pre-stimulus activity across mice revealed an increase in activity before miss trials (number of significant pixels, p=0.005; size of largest cluster, p=0.006) which was largest in somatosensory and visual cortices. **B**, Activity evoked by eye movements had similar spatial structure to the miss-hit pre-stimulus activity maps of some mice (compare with A left panel).

**Figure S10.**
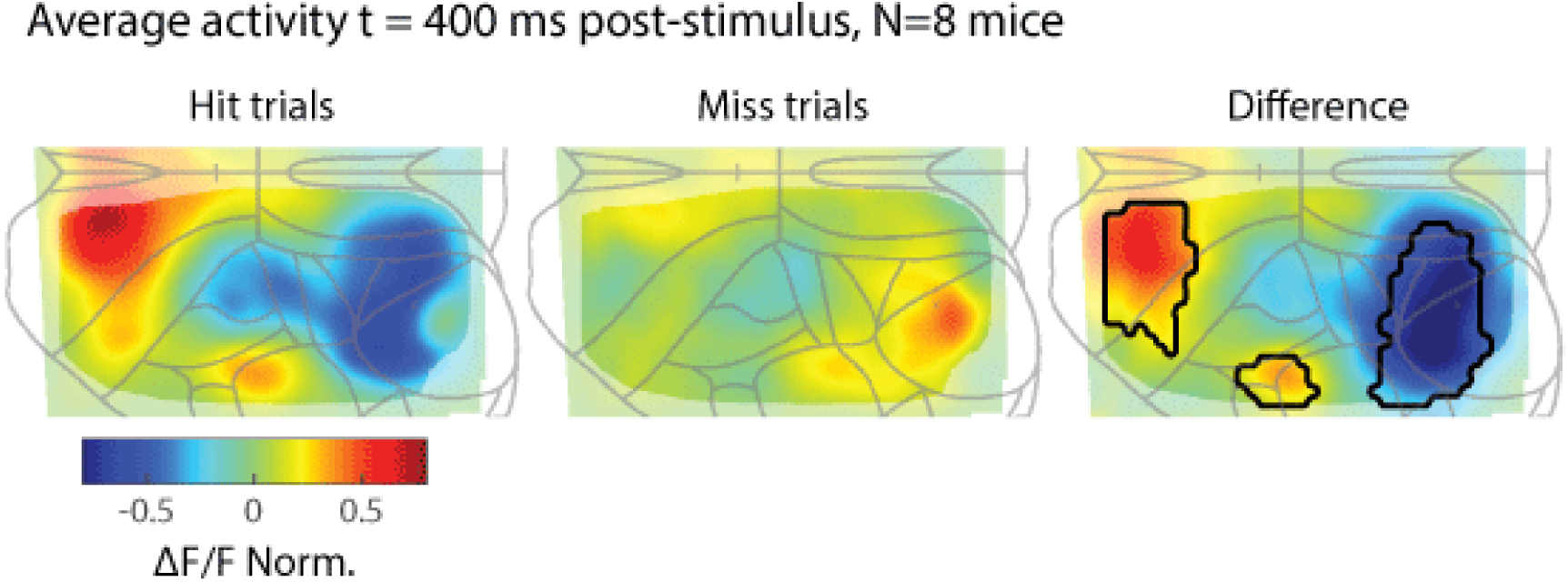
We calculated hit vs. miss activity without subtracting pre-stimulus activity. Left panel: average hit trial activity 400 ms post-stimulus. Middle panel: miss trial activity for same sessions as left panel. Right panel: activity difference between hit and miss. Some differences between this activity difference map and the difference in hit-miss responses (Fig. 3A) are apparent. A significant increase in MOs is preserved, suggesting that overall activity is higher on hit trials. However, we no longer observe a significant increase in parietal cortex. This suggests that the response increase in parietal cortex on hit trials (Fig. 3Avi) is mediated by pre-stimulus noise suppression rather than gain. ΔF/F has been normalized by the maximum pixel value in the hit trial activity map of each mouse to control for overall fluorescence differences between mice. Black outlines in right panel denote clusters of significant pixels (each pixel p<0.05).

### Neural activity in MOs is correlated with response time

The time course of activity in MOs should predict lick response time if this area is important for goal-oriented licking. We sorted single-trial MOs activity by response time to inspect whether the rise of activity was correlated (Fig. S11A). As expected, more delayed response times were predicted by longer times to reach an arbitrary threshold (T_thresh_, z-score 1.5) in MOs after target presentation. T_thresh_ in MOs was often smaller than the lick response time, consistent with this area causing lick behavior. In order to understand how activity in other regions might correlate with response time, we calculated a pixel-wise response time correlation across dorsal cortex (Fig. S11B). We observed a correlation structure similar to the lick rate correlation map (Fig. 4C). MOs and SSp mouth/nose regions were the most highly correlated with some mice also having high correlation in or near SSp-ll (Fig. S11B, evident in grand average). To test if different regions had significantly different response time correlations, we compared three areas of interest: VISp, SSp-ll, and MOs. We observed high variability from mouse to mouse when averaging pixel-wise correlations based on region alone. Inspection of maps for individual mice revealed that clusters of high response time correlation could be highly local. We therefore calculated the center of mass (after rectification) within each region of interest and compared correlations (Fig. S11C). MOs had the highest correlation (0.49 +/- 0.15, p=3.0×10^-6^) and was statistically different from SSp-ll and VISp (p=0.0061 and p=1.1×10^-5^, respectively, paired t-tests). VISp had the lowest correlation, with SSp-ll in between.

**Figure S11.**
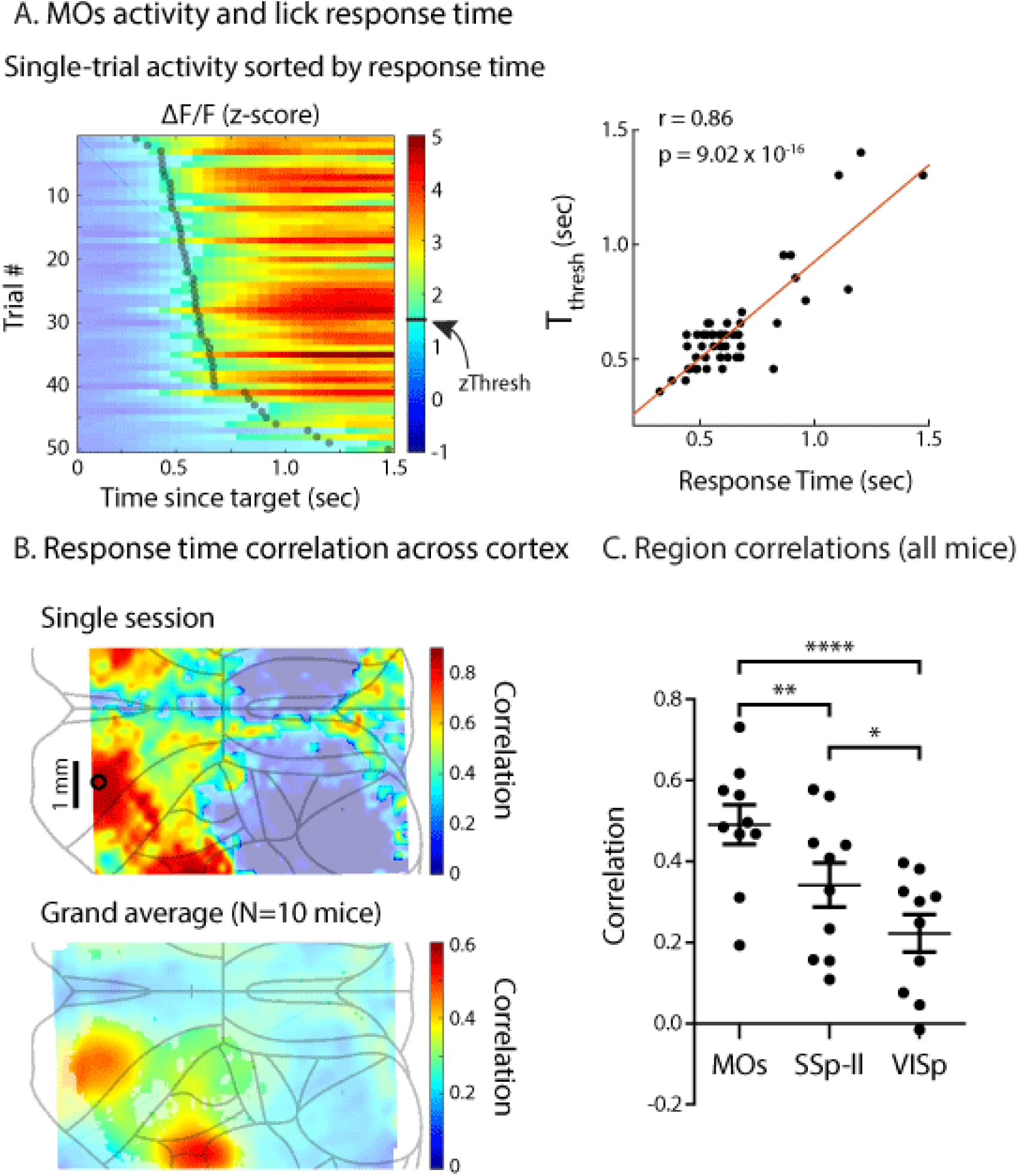
Neural activity in MOs is correlated with response time. **A**, Left: single-trial activity in MOs sorted by response time (transparent green dots) for an example session. Colors become opaque once an arbitrary threshold was crossed (zThresh). Note later lick responses are associated with late threshold crossing. Right: Scatterplot and linear fit of response time and time to reach threshold (T_thresh_). **B**, Response time correlation across cortex for an example session (top) and averaged across all mice (bottom). Black circle in top panel denotes pixel used in A. Opaque pixels have response time correlations above chance (p<0.05) within session (top) and across 10 mice (bottom). **C**, Response time correlation for three regions for all mice. Paired t-test comparisons: MOs and VISp: p=1.11×10^-5^; MOs and SSp-ll: p=0.0061; SSp-ll and VISp: p=0.0195.

**Figure S12.**
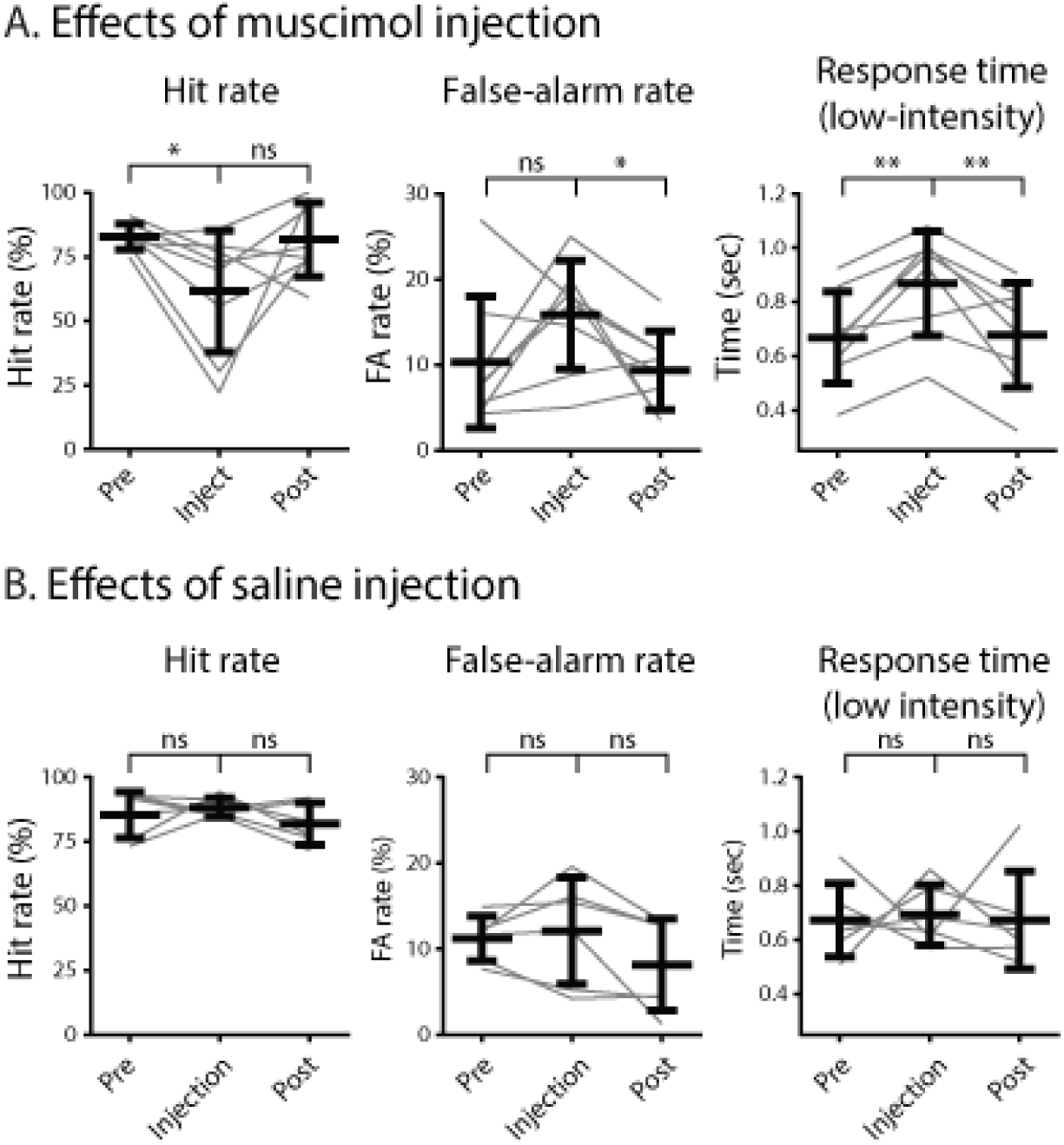
Task performance is not significantly different after saline injection. **A**, Effects of unilateral muscimol injection (n=8 mice; presented here again from Fig. 4E for ease of comparison). Unilateral inactivation was performed in order to monitor the spatial extent of suppression. **B**, Effects of saline injection (n=6 mice). Note the small effects compared to muscimol-injected mice.

## Supplemental Movie

Movie of the face and dorsal cortical activity of a mouse performing the visual detection task. On the left are indicators of time of presentation of a visual stimulus (target), an auditory stimulus (distractor), the lick responses of the animal (lick) and the delivery of water rewards (reward). Note the relationship between activity in Visp with visual stimulation and eye movements, and activity in MOs with lick responses. The colored square to the lower left of the brain activity movie indicates walking speed. Blue is stationary and hotter colors are faster walking.

## References

Allen, William E., Isaac V. Kauvar, Michael Z. Chen, Ethan B. Richman, Samuel J. Yang, Ken Chan, Viviana Gradinaru, Benjamin E. Deverman, Liqun Luo, and Karl Deisseroth. 2017. “Global Representations of Goal-Directed Behavior in Distinct Cell Types of Mouse Neocortex.” Neuron 94 (4): 891–907.e6.

Bisley, James W., and Michael E. Goldberg. 2003. “Neuronal Activity in the Lateral Intraparietal Area and Spatial Attention.” Science 299 (5603): 81–86.

Britten, K. H., W. T. Newsome, M. N. Shadlen, S. Celebrini, and J. A. Movshon. 1996. “A Relationship between Behavioral Choice and the Visual Responses of Neurons in Macaque MT.” Visual Neuroscience 13 (1): 87–100.

Chen, Tsai-Wen, Nuo Li, Kayvon Daie, and Karel Svoboda. 2017. “A Map of Anticipatory Activity in Mouse Motor Cortex.” Neuron. https://doi.org/10.1016/j.neuron.2017.05.005.

Drew, Patrick J., Aaron T. Winder, and Qingguang Zhang. 2018. “Twitches, Blinks, and Fidgets: Important Generators of Ongoing Neural Activity.” The Neuroscientist: A Review Journal Bringing Neurobiology, Neurology and Psychiatry, October, 1073858418805427.

Eggermann, Emmanuel, Yves Kremer, Sylvain Crochet, and Carl C. H. Petersen. 2014. “Cholinergic Signals in Mouse Barrel Cortex during Active Whisker Sensing.” Cell Reports 9 (5): 1654–60.

Erlich, Jeffrey C., Max Bialek, and Carlos D. Brody. 2011. “A Cortical Substrate for Memory-Guided Orienting in the Rat.” Neuron 72 (2): 330–43.

Felleman, D. J., and D. C. Van Essen. 1991. “Distributed Hierarchical Processing in the Primate Cerebral Cortex.” Cerebral Cortex. https://doi.org/10.1093/cercor/1.1.1.

Georgopoulos, A. P., R. E. Kettner, and A. B. Schwartz. 1988. “Primate Motor Cortex and Free Arm Movements to Visual Targets in Three-Dimensional Space. II. Coding of the Direction of Movement by a Neuronal Population.” The Journal of Neuroscience: The Official Journal of the Society for Neuroscience 8 (8): 2928–37.

Goard, Michael J., Gerald N. Pho, Jonathan Woodson, and Mriganka Sur. 2016. “Distinct Roles of Visual, Parietal, and Frontal Motor Cortices in Memory-Guided Sensorimotor Decisions.” eLife. https://doi.org/10.7554/elife.13764.

Guo, Zengcai V., Nuo Li, Daniel Huber, Eran Ophir, Diego Gutnisky, Jonathan T. Ting, Guoping Feng, and Karel Svoboda. 2014. “Flow of Cortical Activity Underlying a Tactile Decision in Mice.” Neuron 81 (1): 179–94.

Hangya, Balázs, Hyun-Jae Pi, Duda Kvitsiani, Sachin P. Ranade, and Adam Kepecs. 2014. “From Circuit Motifs to Computations: Mapping the Behavioral Repertoire of Cortical Interneurons.” Current Opinion in Neurobiology 26 (June): 117–24.

Hanks, Timothy D., and Christopher Summerfield. 2017. “Perceptual Decision Making in Rodents, Monkeys, and Humans.” Neuron 93 (1): 15–31.

Hernández, Adrián, Verónica Nácher, Rogelio Luna, Antonio Zainos, Luis Lemus, Manuel Alvarez, Yuriria Vázquez, Liliana Camarillo, and Ranulfo Romo. 2010. “Decoding a Perceptual Decision Process across Cortex.” Neuron 66 (2): 300–314.

Inagaki, Hidehiko K., Lorenzo Fontolan, Sandro Romani, and Karel Svoboda. 2019. “Discrete Attractor Dynamics Underlies Persistent Activity in the Frontal Cortex.” Nature 566 (7743): 212–17.

Inagaki, Hidehiko K., Miho Inagaki, Sandro Romani, and Karel Svoboda. 2018. “Low-Dimensional and Monotonic Preparatory Activity in Mouse Anterior Lateral Motor Cortex.” The Journal of Neuroscience: The Official Journal of the Society for Neuroscience 38 (17): 4163–85.

Kawai, Risa, Timothy Markman, Rajesh Poddar, Raymond Ko, Antoniu L. Fantana, Ashesh K. Dhawale, Adam R. Kampff, and Bence P. Ölveczky. 2015. “Motor Cortex Is Required for Learning but Not for Executing a Motor Skill.” Neuron 86 (3): 800–812.

Kebschull, Justus M., Pedro Garcia da Silva, Ashlan P. Reid, Ian D. Peikon, Dinu F. Albeanu, and Anthony M. Zador. 2016. “High-Throughput Mapping of Single-Neuron Projections by Sequencing of Barcoded RNA.” Neuron 91 (5): 975–87.

Keller, Georg B., and Thomas D. Mrsic-Flogel. 2018. “Predictive Processing: A Canonical Cortical Computation.” Neuron 100 (2): 424–35.

Khan, Adil G., and Sonja B. Hofer. 2018. “Contextual Signals in Visual Cortex.” Current Opinion in Neurobiology 52 (October): 131–38.

Komiyama, Takaki, Takashi R. Sato, Daniel H. O’Connor, Ying-Xin Zhang, Daniel Huber, Bryan M. Hooks, Mariano Gabitto, and Karel Svoboda. 2010. “Learning-Related Fine-Scale Specificity Imaged in Motor Cortex Circuits of Behaving Mice.” Nature 464 (7292): 1182– 86.

Kyriakatos, Alexandros, Vijay Sadashivaiah, Yifei Zhang, Alessandro Motta, Matthieu Auffret, and Carl C. H. Petersen. 2016. “Voltage-Sensitive Dye Imaging of Mouse Neocortex during a Whisker Detection Task.” Neurophotonics. https://doi.org/10.1117/1.nph.4.3.031204.

Lafuente, Victor de, and Ranulfo Romo. 2006. “Neural Correlate of Subjective Sensory Experience Gradually Builds up across Cortical Areas.” Proceedings of the National Academy of Sciences of the United States of America 103 (39): 14266–71.

Lee, A. Moses, Jennifer L. Hoy, Antonello Bonci, Linda Wilbrecht, Michael P. Stryker, and Cristopher M. Niell. 2014. “Identification of a Brainstem Circuit Regulating Visual Cortical State in Parallel with Locomotion.” Neuron 83 (2): 455–66.

Lee, Seung-Hee, and Yang Dan. 2012. “Neuromodulation of Brain States.” Neuron 76 (1): 209– 22.

Lee, Soohyun, Illya Kruglikov, Z. Josh Huang, Gord Fishell, and Bernardo Rudy. 2013. “A Disinhibitory Circuit Mediates Motor Integration in the Somatosensory Cortex.” Nature Neuroscience 16 (11): 1662–70.

Leinweber, Marcus, Daniel R. Ward, Jan M. Sobczak, Alexander Attinger, and Georg B. Keller. 2017. “A Sensorimotor Circuit in Mouse Cortex for Visual Flow Predictions.” Neuron 96 (5): 1204.

Li, Nuo, Tsai-Wen Chen, Zengcai V. Guo, Charles R. Gerfen, and Karel Svoboda. 2015. “A Motor Cortex Circuit for Motor Planning and Movement.” Nature 519 (7541): 51–56.

Li, Nuo, Kayvon Daie, Karel Svoboda, and Shaul Druckmann. 2016. “Robust Neuronal Dynamics in Premotor Cortex during Motor Planning.” Nature 532 (7600): 459–64.

Madisen, Linda, Aleena R. Garner, Daisuke Shimaoka, Amy S. Chuong, Nathan C. Klapoetke, Lu Li, Alexander van der Bourg, et al. 2015. “Transgenic Mice for Intersectional Targeting of Neural Sensors and Effectors with High Specificity and Performance.” Neuron 85 (5): 942–58.

Makino, Hiroshi, Chi Ren, Haixin Liu, An Na Kim, Neehar Kondapaneni, Xin Liu, Duygu Kuzum, and Takaki Komiyama. 2017. “Transformation of Cortex-Wide Emergent Properties during Motor Learning.” Neuron 94 (4): 880–90.e8.

Ma, Ying, Mohammed A. Shaik, Sharon H. Kim, Mariel G. Kozberg, David N. Thibodeaux, Hanzhi T. Zhao, Hang Yu, and Elizabeth M. C. Hillman. 2016. “Wide-Field Optical Mapping of Neural Activity and Brain Haemodynamics: Considerations and Novel Approaches.” Philosophical Transactions of the Royal Society of London. Series B, Biological Sciences 371 (1705). https://doi.org/10.1098/rstb.2015.0360.

McCormick, D. A. 1992. “Neurotransmitter Actions in the Thalamus and Cerebral Cortex and Their Role in Neuromodulation of Thalamocortical Activity.” Progress in Neurobiology 39 (4): 337–88.

McCormick, David A., Matthew J. McGinley, and David B. Salkoff. 2015. “Brain State Dependent Activity in the Cortex and Thalamus.” Current Opinion in Neurobiology 31 (April): 133–40.

McGinley, Matthew J., Stephen V. David, and David A. McCormick. 2015. “Cortical Membrane Potential Signature of Optimal States for Sensory Signal Detection.” Neuron 87 (1): 179– 92.

McGinley, Matthew J., Martin Vinck, Jacob Reimer, Renata Batista-Brito, Edward Zagha, Cathryn R. Cadwell, Andreas S. Tolias, Jessica A. Cardin, and David A. McCormick. 2015. “Waking State: Rapid Variations Modulate Neural and Behavioral Responses.” Neuron 87 (6): 1143–61.

Mitra, Anish, Andrew Kraft, Patrick Wright, Benjamin Acland, Abraham Z. Snyder, Zachary Rosenthal, Leah Czerniewski, et al. 2018. “Spontaneous Infra-Slow Brain Activity Has Unique Spatiotemporal Dynamics and Laminar Structure.” Neuron 98 (2): 297–305.e6.

Muñoz, William, and Bernardo Rudy. 2014. “Spatiotemporal Specificity in Cholinergic Control of Neocortical Function.” Current Opinion in Neurobiology 26 (June): 149–60.

Musall, Simon, Matthew T. Kaufman, Ashley L. Juavinett, Steven Gluf, and Anne K. Churchland. 2019. “Single-Trial Neural Dynamics Are Dominated by Richly Varied Movements.” https://doi.org/10.1101/308288.

Nelson, Anders, and Richard Mooney. 2016. “The Basal Forebrain and Motor Cortex Provide Convergent yet Distinct Movement-Related Inputs to the Auditory Cortex.” Neuron. https://doi.org/10.1016/j.neuron.2016.03.031.

Nelson, Anders, David M. Schneider, Jun Takatoh, Katsuyasu Sakurai, Fan Wang, and Richard Mooney. 2013. “A Circuit for Motor Cortical Modulation of Auditory Cortical Activity.” The Journal of Neuroscience: The Official Journal of the Society for Neuroscience 33 (36): 14342–53.

Nichols, Thomas E., and Andrew P. Holmes. 2002. “Nonparametric Permutation Tests for Functional Neuroimaging: A Primer with Examples.” Human Brain Mapping 15 (1): 1–25.

Niell, Cristopher M., and Michael P. Stryker. 2010. “Modulation of Visual Responses by Behavioral State in Mouse Visual Cortex.” Neuron 65 (4): 472–79.

Oh, Seung Wook, Julie A. Harris, Lydia Ng, Brent Winslow, Nicholas Cain, Stefan Mihalas, Quanxin Wang, et al. 2014. “A Mesoscale Connectome of the Mouse Brain.” Nature 508 (7495): 207–14.

Petersen, Carl C. H. 2007. “The Functional Organization of the Barrel Cortex.” Neuron 56 (2): 339–55.

Pi, Hyun-Jae, Balázs Hangya, Duda Kvitsiani, Joshua I. Sanders, Z. Josh Huang, and Adam Kepecs. 2013. “Cortical Interneurons That Specialize in Disinhibitory Control.” Nature 503 (7477): 521–24.

Pinto, Lucas, Michael J. Goard, Daniel Estandian, Min Xu, Alex C. Kwan, Seung-Hee Lee, Thomas C. Harrison, Guoping Feng, and Yang Dan. 2013. “Fast Modulation of Visual Perception by Basal Forebrain Cholinergic Neurons.” Nature Neuroscience 16 (12): 1857–63.

Poulet, James F. A., and Sylvain Crochet. 2018. “The Cortical States of Wakefulness.” Frontiers in Systems Neuroscience 12: 64.

Reimer, Jacob, Matthew J. McGinley, Yang Liu, Charles Rodenkirch, Qi Wang, David A. McCormick, and Andreas S. Tolias. 2016. “Pupil Fluctuations Track Rapid Changes in Adrenergic and Cholinergic Activity in Cortex.” Nature Communications 7 (November): 13289.

Robinson, D. L., E. M. Bowman, and C. Kertzman. 1995. “Covert Orienting of Attention in Macaques. II. Contributions of Parietal Cortex.” Journal of Neurophysiology 74 (2): 698– 712.

Schneider, David M., and Richard Mooney. 2018. “How Movement Modulates Hearing.” Annual Review of Neuroscience 41 (July): 553–72.

Schneider, David M., Anders Nelson, and Richard Mooney. 2014. “A Synaptic and Circuit Basis for Corollary Discharge in the Auditory Cortex.” Nature 513 (7517): 189–94.

Schwarz, Lindsay A., Kazunari Miyamichi, Xiaojing J. Gao, Kevin T. Beier, Brandon Weissbourd, Katherine E. DeLoach, Jing Ren, et al. 2015. “Viral-Genetic Tracing of the Input-Output Organization of a Central Noradrenaline Circuit.” Nature 524 (7563): 88–92.

Shimaoka, Daisuke, Kenneth D. Harris, and Matteo Carandini. 2018. “Effects of Arousal on Mouse Sensory Cortex Depend on Modality.” Cell Reports 25 (11): 3230.

Siegel, Markus, Timothy J. Buschman, and Earl K. Miller. 2015. “Cortical Information Flow during Flexible Sensorimotor Decisions.” Science 348 (6241): 1352–55.

Steinmetz, Nicholas A., Christina Buetfering, Jerome Lecoq, Christian R. Lee, Andrew J. Peters, Elina A. K. Jacobs, Philip Coen, et al. 2017. “Aberrant Cortical Activity in Multiple GCaMP6-Expressing Transgenic Mouse Lines.” eNeuro 4 (5). https://doi.org/10.1523/ENEURO.0207-17.2017.

Stringer, Carsen, Marius Pachitariu, Nicholas Steinmetz, Charu Bai Reddy, Matteo Carandini, and Kenneth D. Harris. 2019. “Spontaneous Behaviors Drive Multidimensional, Brainwide Activity.” Science 364 (6437): 255.

Swets, John A., and David M. Green. 1978. “Applications of Signal Detection Theory.” Psychology: From Research to Practice. https://doi.org/10.1007/978-1-4684-2487-4_19.

Travers, J. B., L. A. Dinardo, and H. Karimnamazi. 1997. “Motor and Premotor Mechanisms of Licking.” Neuroscience and Biobehavioral Reviews 21 (5): 631–47.

Wekselblatt, Joseph B., Erik D. Flister, Denise M. Piscopo, and Cristopher M. Niell. 2016. “Large-Scale Imaging of Cortical Dynamics during Sensory Perception and Behavior.” Journal of Neurophysiology 115 (6): 2852–66.

Zagha, Edward, Amanda E. Casale, Robert N. S. Sachdev, Matthew J. McGinley, and David A. McCormick. 2013. “Motor Cortex Feedback Influences Sensory Processing by Modulating Network State.” Neuron 79 (3): 567–78.

Zagha, Edward, Xinxin Ge, and David A. McCormick. 2015. “Competing Neural Ensembles in Motor Cortex Gate Goal-Directed Motor Output.” Neuron 88 (3): 565–77.

Zagha, Edward, and David A. McCormick. 2014. “Neural Control of Brain State.” Current Opinion in Neurobiology 29 (December): 178–86.

Zhang, Siyu, Min Xu, Tsukasa Kamigaki, Johnny Phong Hoang Do, Wei-Cheng Chang, Sean Jenvay, Kazunari Miyamichi, Liqun Luo, and Yang Dan. 2014. “Selective Attention. Long-Range and Local Circuits for Top-down Modulation of Visual Cortex Processing.” Science 345 (6197): 660–65.

## References

Chen, Tsai-Wen, Trevor J. Wardill, Yi Sun, Stefan R. Pulver, Sabine L. Renninger, Amy Baohan, Eric R. Schreiter, et al. 2013. “Ultrasensitive Fluorescent Proteins for Imaging Neuronal Activity.” Nature 499 (7458): 295–300.

Massimini, Marcello, Reto Huber, Fabio Ferrarelli, Sean Hill, and Giulio Tononi. 2004. “The Sleep Slow Oscillation as a Traveling Wave.” The Journal of Neuroscience: The Official Journal of the Society for Neuroscience 24 (31): 6862–70.

Otchy, Timothy M., Steffen B. E. Wolff, Juliana Y. Rhee, Cengiz Pehlevan, Risa Kawai, Alexandre Kempf, Sharon M. H. Gobes, and Bence P. Ölveczky. 2015. “Acute off-Target Effects of Neural Circuit Manipulations.” Nature 528 (7582): 358–63.

Sirotin, Yevgeniy B., and Aniruddha Das. 2009. “Anticipatory Haemodynamic Signals in Sensory Cortex Not Predicted by Local Neuronal Activity.” Nature 457 (7228): 475–79.

Steriade, M., A. Nunez, and F. Amzica. 1993. “A Novel Slow (< 1 Hz) Oscillation of Neocortical Neurons in Vivo: Depolarizing and Hyperpolarizing Components.” The Journal of Neuroscience. https://doi.org/10.1523/jneurosci.13-08-03252.1993.

